# Modulation of Calcium Signaling and Metabolic Pathways in Endothelial Cells with Magnetic Fields

**DOI:** 10.1101/2023.10.07.561321

**Authors:** Oksana Gorobets, Svitlana Gorobets, Tatyana Polyakova, Vitalii Zablotskii

## Abstract

Calcium signaling plays a crucial role in various physiological processes, including muscle contraction, cell division, and neurotransmitter release. Dysregulation of calcium levels and signaling has been linked to a range of pathological conditions such as neurodegenerative disorders, cardiovascular disease, and cancer. Here, we suggest that in the endothelium, calcium ion channel activity and calcium signaling can be modulated by applying either a time-varying or static gradient magnetic field (MF). This modulation is achieved by exerting magnetic forces or torques on either biogenic or non-biogenic magnetic nanoparticles that are bound to endothelial cell membranes. Since calcium signaling in endothelial cells induces neuromodulation and influences blood flow control, treatment with a magnetic field shows promise for regulating neurovascular coupling and treating vascular dysfunctions associated with aging and neurodegenerative disorders. Furthermore, magnetic treatment can enable control over the decoding of Ca signals, ultimately impacting protein synthesis. The ability to modulate calcium wave frequencies using MFs and the MF-controlled decoding of Ca signaling present promising avenues for treating diseases characterized by calcium dysregulation.

## Introduction

The ability to process information at the cellular and subcellular levels is fundamental to important processes in living organisms - from cell fertilization and division to short-term memory, aging, and the development of many diseases. Biological information transfer and control of processes in living organisms occur through physiological oscillations, which can take the form of individual spikes, periodic oscillations, and propagating waves. The mechanisms underlying such physiological oscillations and their roles remain largely unknown [1]. A vast amount of experimental evidence has accumulated regarding calcium oscillations, which demonstrate the crucial role of calcium ion signals in all levels of structural organization of the organism, from subcellular to large-scale calcium spikes and waves in neurons of the brain [2] and long-range calcium signals in plants [3,4]. Calcium signals, which propagate as concentration waves of calcium, direct the functioning of the entire organism, starting from the moment of the emergence of life when sperm introduce phospholipase C-ζ into the egg and a slow oscillating wave of Ca^2+^ moves through the egg cell, triggering fertilization [5,6], and ending with its extinction, accompanied by calcium necrotic waves that contribute to the death of the organism [7]. During the development and life of an organism, biological information is transmitted through frequency-modulated waves of calcium, similar to radio signals [5]. This information is encoded through a complex set of channels and transporters using specific frequency modulation of calcium waves, and then transmitted to receivers, where it is decoded by several sensors and effectors, resulting in a conversion into specific cellular processes and metabolic pathways.

Despite extensive knowledge accumulated about the mechanisms of information processing transmitted by calcium signals, the principles of its encoding and decoding remain largely unclear. It seems that in most cases, the transmission of biological information follows a principle exemplified by the activation of Ca^2+^/calmodulin-dependent protein kinase II (CaMKII), an enzymatic complex consisting of dimers of hexameric rings. CaMKII is particularly sensitive to the frequency of intracellular Ca^2+^ oscillations, where only high-frequency oscillations provide sequential and autonomous activation of individual catalytic domains, and the duration or amplitude of the pulse does not affect its activation [8]. The process of decoding is based on the kinetics of calcium binding and unbinding to kinases and phosphatases, which, respectively, activate and deactivate target proteins [5]. Each molecule involved in decoding Ca^2+^ oscillations can recognize oscillatory patterns within a specific frequency range. Different decoding proteins have their own specific frequency ranges (with only a small overlap between them), indicating their specific roles in cells, such as activating specific cellular programs. Therefore, different frequency ranges correspond to different Ca^2+^ binding proteins that control different metabolic processes and generate different cellular signals. The frequencies of intracellular Ca^2+^ oscillations are specific to each cell type and range from about 1 mHz in endothelial cells to about 1000 mHz in cardiac cells and neurons [5,9]. The frequency of calcium oscillations determines the pace of the body’s biological clocks. In neuronal networks, the frequency of Ca-mediated spike oscillations plays a crucial role in determining biological timing and organisms’ perception of time. Spike oscillations in neuronal circuits are tightly regulated and synchronized, facilitating various cognitive processes such as attention, memory, and decision-making [10,11].

As oscillating calcium signals are associated with an individual’s cognitive abilities and memory, and magnetic fields (MFs) are capable of affecting the dynamic patterns of calcium, such as releasing calcium from intracellular depots and increasing its concentration in the cytosol [12–15], the influence of magnetic fields on cognitive processes such as improving attention, memory, and decision-making abilities [16–18] no longer seem as unexpected as they might at first glance. The transmission of calcium signals is a crucial component of brain plasticity and is involved in various forms of plasticity, including synaptic plasticity, neurogenesis, and dendritic branching [19]. In the heart, calcium oscillations play a critical role in regulating heart rate and rhythm [20]. The treatment of human umbilical vein endothelial cells with a rotating magnetic field (4 Hz and0.2 T) demonstrated an increase in intracellular Ca^2+^ concentration. When *Caenorhabditis elegans* was exposed to this magnetic field for a prolonged period (hours), it surprisingly resulted in an extended lifespan [21]. Overall, calcium oscillations represent a universal and fundamental signaling mechanism that plays a decisive role in various physiological processes in living beings.

Disruptions in Ca signaling can lead to the development of various diseases such as osteoporosis, arterial hypertension, cardiac arrhythmia, diabetes, and neurological disorders including Alzheimer’s and Parkinson’s diseases. By properly modulating the frequency of Ca signal transmission, specific pathways and processes involved in the development or progression of many diseases can be pharmacologically targeted. For example, drugs that increase or decrease the frequency of Ca oscillations can be used to target specific types of cancer cells [22]. In addition, modulating Ca signal transmission can help reduce inflammation in autoimmune diseases by regulating the function of immune cells [23]. However, to date, there is no universal mechanism or physical tool that allows for selective control of calcium ion channels, thereby modulating calcium oscillations and waves to achieve therapeutic effects. The human genome encodes at least 400 members of ion channel families (∼1.5%), making it the second-largest class of membrane proteins for drug design after G protein-coupled receptors (GPCRs). About 18% of the small molecule drugs listed in the ChEMBL database [24] target ion channels [25]. Although ion channels are widely recognized as the basis of many diseases (including cancers, dementia, diabetes, and asthma), approved drugs are available for only a small percentage of this protein class (approximately 8%), despite concerted drug discovery efforts over the past 30 years [25]. Over the past decades, ion channels have been considered challenging targets for drugs due to the difficulty in achieving drug selectivity and specificity. Given that ion channels are present in nearly all living cells, and a significant number share structural similarities [26] [27], the primary focus of numerous researchers lies in discovering a unified method enabling selective control of ion channel activity. Hence, the development of physical methods for targeting ion channels stands as an alternative to existing drugs like small molecules and antibodies. In this study, we propose the application of magnetic fields and magnetic nanoparticles as a highly selective and specific method for targeting calcium ion channels. This innovative approach enables precise targeting of particular organs, tissues, and cells. Furthermore, we propose a non-invasive method utilizing magnetic fields to modulate calcium signal oscillations.

Through theoretical modeling and computer simulations, we found magnetic field regimes that provide frequency or amplitude modulation of ion channel activity and waves of free calcium concentration. We show how with suitable tuning of calcium oscillation frequencies, it is possible to selectively control the decoding protein functions and the synthesis of related proteins. These results expand our understanding of the mechanisms of ion channel activation and inactivation and can serve as a reliable theoretical basis for developing new approaches to treating a wide range of human diseases.

## Results

### Model and mechanisms of MF impacts on ion channels

We now construct a mathematical model of the MF influence on the key intracellular and extracellular processes incorporated in Ca^2+^ dynamics. We start with the robust models of Plank et al. (2006) [28] and Wiesner et al. (1996) [29], which have been verified by numerous experiments on Ca^2+^ dynamics [30–34].The basic equations of our model are given in the Section 4 (Method). Our generalization of the model [28] takes into account the following key factors related to a MF. The first, the existence of biogenic magnetic nanoparticles (BMNs) practically in all organs of organisms [35–41], Fig. 1. The second, in an organism, the BMNs self-assemble into chains mainly located on lipid membranes of endothelial [42], cardiac [43] and other cells [44]. The third, many types of membrane ion channels are sensitive to small shear stress which can gate these channels [45]. Finally, when an external magnetic field is applied, it causes mechanical forces to act on BMN chains. These mechanical forces are then transmitted into the cell membrane, causing shear stress within it. Sources of BMNs are: aging (as the body ages, mitochondrial dysfunction in cells increases iron accumulation in various tissues and causes iron-dependent brain degeneration [46]), hypoxia, environmental contamination [47,48], magnetic nanoparticles injected as contrast agent at MRI [49,50], etc. It should be noted that mechanical shear stress in a cell membrane directly affects calcium influx, q_in_ as described in model [28] by Eq. 8 in the Method Section.

**Fig. 1.**
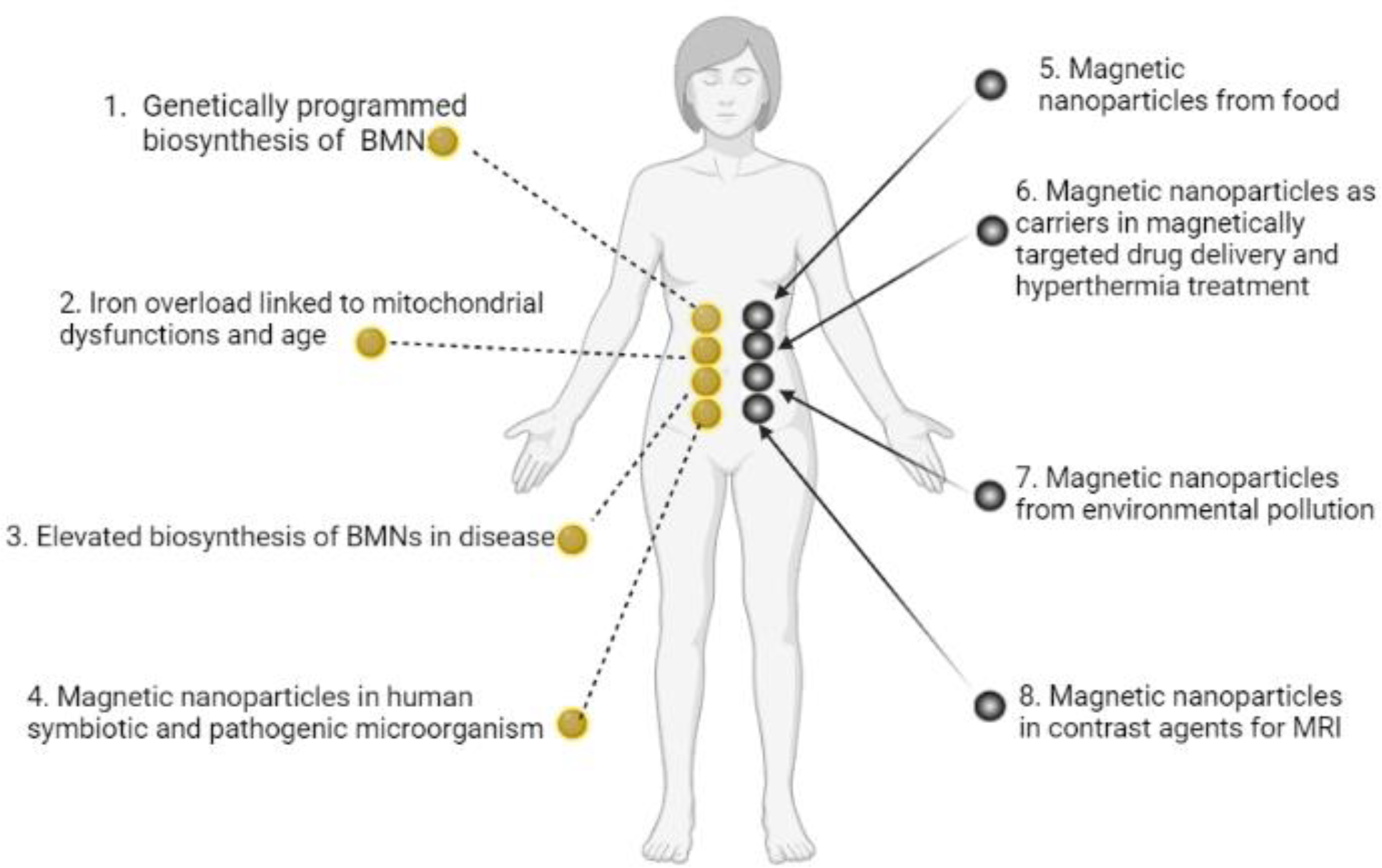
Origins of BMNs and other magnetic nanoparticles in an organism. 1. Genetically programmed biosynthesis of chains of biogenic magnetic nanoparticles [37,51–55], see also the review paper [41]. 2. Iron overload linked to mitochondrial dysfunctions and age-related diseases [46] 3. Elevated biosynthesis of biogenic magnetic nanoparticles in disease (a number of neurodegenerative diseases and cancer) [36,56–58], see also the review paper [41]. 4. Magnetic nanoparticles in human symbiotic and pathogenic microorganism [59,60]. 5. Magnetic nanoparticles from food (plant [61]), mushroom [62], animal [63]). 6. Magnetic nanoparticles as carriers in magnetically targeted drug and cell delivery (see the review papers [64–68]) and hyperthermia [69,70]. 7. Magnetic nanoparticles from environment pollution [47,48]. For example, airborne magnetite pollution particles < ∼200 nm in size can access the brain directly via the olfactory and/or trigeminal nerves, bypassing the blood-brain barrier [48]. 8. Magnetic nanoparticles in contrast agents for MRI [49] [50]. In particular, BMNs are experimentally revealed in human brain [35,36], heart [37], liver [37], spleen [37], ethmoid bone [38], adrenal glands [39]. BMNs in animals and human are chemically pure magnetite or maghemite and was found to be organized into magnetically interacting clusters and linear membrane-bound chains [42,71–73] as well as BMNs in magnetotactic bacteria. For example, human brain contains the intracellular magnetite crystals forming chains and are bound to the cell membrane, and have a saturation magnetization Ms = 4.5×10^5^ A/m [74]; there are up to 80 magnetite crystals in a chain in the human brain according to [75]. The chains of BMN are found experimentally in capillary walls that are formed by a single layer of endothelial cells [42]. In pathology, biogenic magnetic nanoparticles are revealed in tumour tissue in high amounts [36,76] at the cell membrane [77].

### How a magnetic field works

Below, we explain how an externally applied magnetic field interacting with BMN chains can influence the key molecular processes that regulate endothelial calcium dynamics. The main processes included in the model of Ca^2+^ dynamics are as follows [28]. 1. The internal production of inositol phosphate signaling molecules (IP3), which include several processes such as the activation of a G-protein and phospholipase C and the cleavage of PIP_2_. 2. The intracellular receptor for IP3 is responsible for generation and control of Ca^2+^ signals. IP3 opens Ca^2+^ channels in the endoplasmic reticulum (ER). 3. Both the rate of IP3 production and the rate of internal Ca^2+^ release can be enhanced by cytosolic free Ca^2+^. 4. The influx rate of Ca^2+^ due to the process termed capacitive calcium entry [78] is an increasing function of the degree of depletion of the ER below resting levels, as well as the Ca^2+^ concentration difference across the plasma membrane. 5. The rate of Ca^2+^ influx through shear-gated channels is an increasing function of the mechanical shear stress to which endothelial cells are exposed. 6. Cytosolic Ca^2+^ is resequestered back into the ER by a Ca^2+^-ATPase and is pumped out of the cell by a plasma membrane Ca^2+-^ATPase and Ca^2+^-Na^+^ exchanger. 7. Cytosolic Ca^2+^ is reversibly buffered to proteins such as calmodulin. Among these processes, the central player is the mechanical shear stress in the cell membrane caused by blood flow. The magnetic force acting on chains of magnetic nanoparticles creates additional shear stress in the membrane, which, together with the blood flow, regulates mechanosensitive calcium channels.

In Section 4 (Method), these processes are mathematically described by the system of differential equations (7-10). The wall shear stress (WSS), which plays a key role in regulating calcium dynamics, is also included in these equations [28,29]. To take into account the effects of magnetic fields in Eqs. (7-10), we incorporate the magnetic-field-induced shear stress in addition to that induced by blood flow as follows

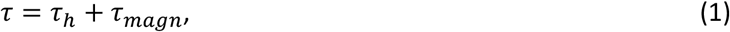

where τ_*h*_ is WSS induced by hydrodynamic blood flow in an endothelial cell (the typical values of WSS of blood flow in vessels are: τ_*h*_ = 1Pa for artery and τ_*h*_ = 0.1 Pa [79] for capillary) and τ_*magn*_ is WSS induced by the magnetic forces acting on the chain of magnetic nanoparticles embedded in cell membrane. In this context, the chains of BMNs located on the membranes of endothelial cells act as effectors, responding to magnetic fields by modulating mechanosensitive ion channels. Further, we consider the WSS induced by gradient and uniform MFs.

Below we show theoretically how to modulate ion channels activity and modulate calcium signaling frequency with uniform and gradient magnetic fields.

### Ca^2+^ ion channel gating by MFs through wall shear stress

To analytically describe the influx of Ca^2+^, we utilize Eqs. 1-3 and the model [28] that investigates how membrane ion channels respond to shear stress. The application of shear stress induces a uniaxial tension gradient in the membrane and deforms it. The deformation of the membrane imparts a strain energy density, *W*, which depends upon the applied shear stress. The fraction of channels in the open state depends on the strain energy density as follows [29]:

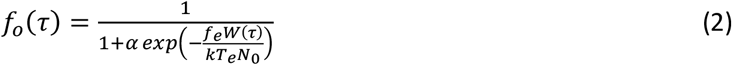

where τ is applied wall shear stress (WSS), *N*_0_ is channel density per unit area of cell membrane, *k* is the Boltzmann constant, *T*_*e*_ is the absolute temperature, and *α* (*α* ≥ 0) is a measure of the probability that a channel is in the open state in the no-load case, namely, (1 + *α*)^−1^ is the probability that a channel is in the open state in the no-load case *W*=0, *f*_*e*_ (0 ≤ *f*_*e*_ ≤ 1) is the fraction of the energy of the membrane that gates the WSS-sensitive Ca^2+^ channels.

The strain energy density function has the form for a two-dimensional membrane [29,80] (Fig. S1, Fig. S2):

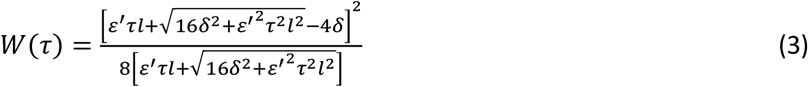

where *ε*^′^ is the fraction of the applied load borne by submembranous structures, *δ* is membrane shear modulus, *l* is cell length in the flow direction.

The shear-dependent Ca-influx is proportional to the fraction of open Ca^2+^ channels in the plasma membrane [29], which has a Boltzmann dependence [81] on the strain energy density in the plasma membrane *W*(τ). This gives a sigmoidal relation between WSS-dependent Ca-influx and the applied WSS. Consequently, the following expression is valid for Ca^2+^ influx [28]:

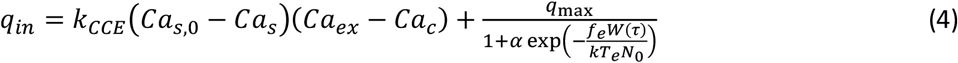

where *Ca*_*s*_ is concentration of Ca^2+^ in the internal stores, *Ca*_*s*,0_ is an initial value of the concentration of Ca^2+^ in the internal stores, *Ca*_*c*_ is concentration of free Ca^2+^ in the cytosol, *Ca*_*ex*_ is concentration of external Ca^2+^, *k*_*CCE*_ and *q*_max_ are the constants. In Eqs. 2 and 4.

Let us calculate the WSS in (2) and (3) induced by gradient magnetic field application. First, we consider a chain of magnetic nanoparticles embedded in a cell membrane under the influence of a gradient magnetic field (Fig. 2A).

**Fig. 2.**
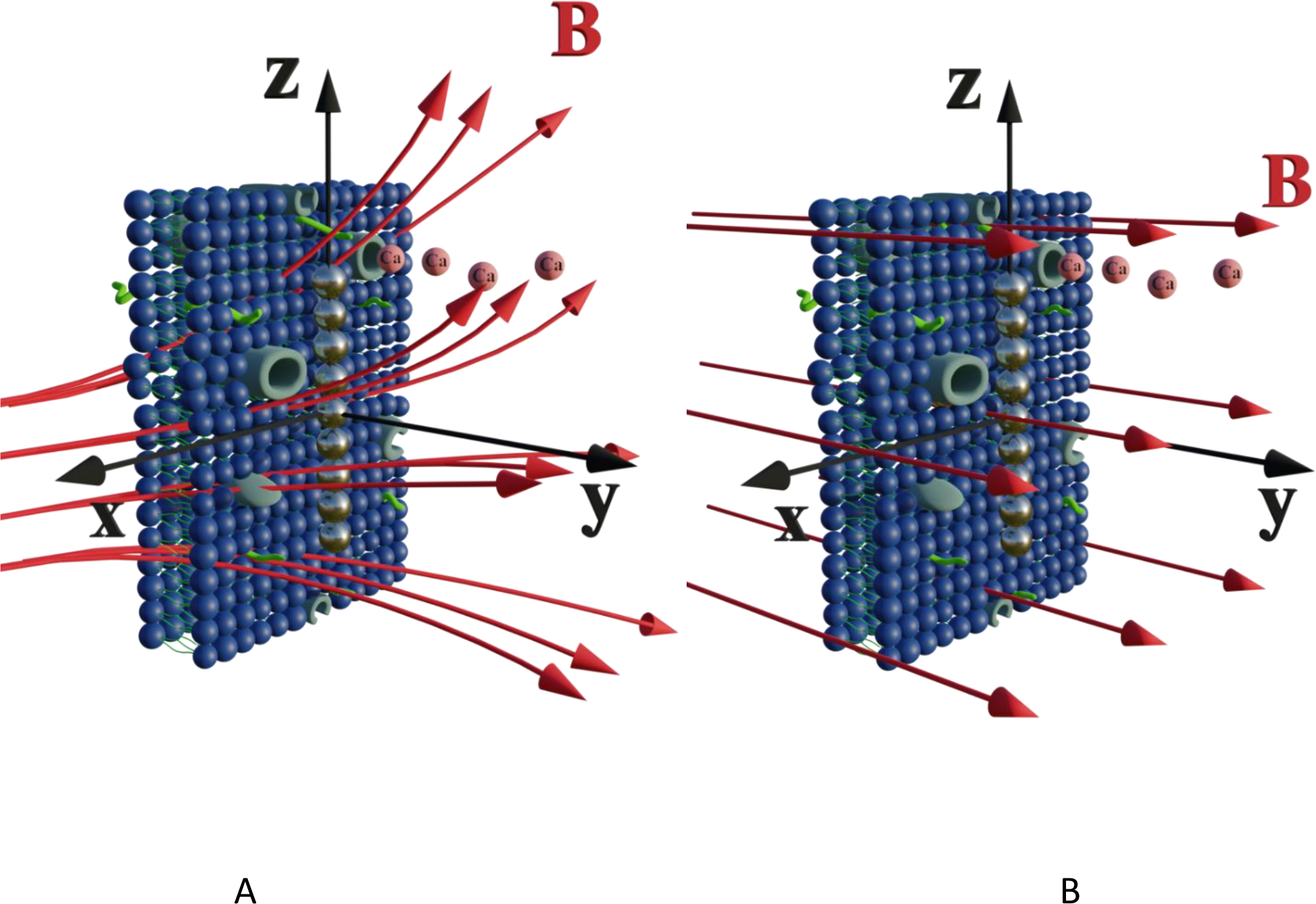
Schematic of a chain of magnetic nanoparticles on a cell membrane under the influence of a) gradient and b) uniform magnetic field. Two calcium ion channels are depicted in the vicinity of the chain. The red arrows are magnetic lines.

The magnetic gradient force acting on a BMN chain is 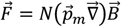, where 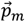 is the magnetic nanoparticle magnetic dipole moment, *N* is the number of magnetic nanoparticles in the chain, 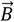 is the magnetic field induction. The WSS is 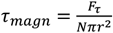 where *F*_τ_ is the in-plane component of magnetic gradient force, *r* is the magnetic nanoparticle radius. Then, in a gradient MF the stress is

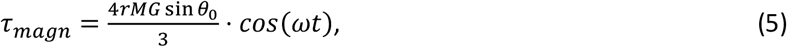

where *M* is magnetization of magnetic nanoparticle (which is a function of applied magnetic field), 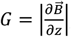, *θ*_0_ is the angle between 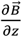 and the normal to the plane of membrane in the case when z-axis is chosen along the direction of 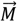 (Fig. 2 A). The parameter *G* characterizes MF gradient and it is considered as an oscillating harmonic function with the frequency *ω*. For a static gradient MF, the share stress is described by Eq.5 at *ω* = 0.

Now we calculate the WSS in (3) induced by uniform magnetic field application. We consider a chain of magnetic nanoparticles embedded in a cell membrane under the influence of a uniform MF (Fig. 2 B).

The magnetic torque can be written as a vector sum of its in-plane and out-of-plane components 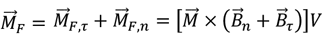,where 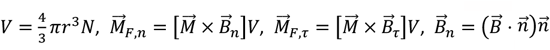 is the out-of-plane component of the magnetic field induction, 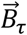 is the in-plane component of magnetic field induction, 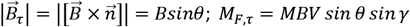is the absolute value of the in-plane component of the magnetic torque producing WSS (Fig. 2 B). Then in the case of uniform magnetic field

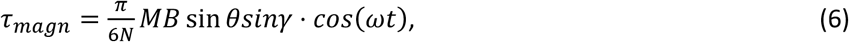

where *θ* is the angle between the normal to the cell membrane and MF direction, γ is the angle between the MF in-plane component and magnetization of the chain of magnetic nanoparticles (Fig. 2 B). In Eq.6, in a general case, the magnetic field flux density is considered as oscillating harmonic function, while for a static MF *ω* = 0.

It should be noted here that even for a low magnetic field (MF) such as the geomagnetic field, the shear stress is not small. For example, an estimate (Eq. 6) gives a magnitude of shear stress of about 1.3 Pa for the geomagnetic field *B*=0.05 mT and *N* = 10, *r* = 100 *nm, M*_*S*_ = 510 kA/m.

Thus, Eqs. 5 and 6 describe magnetic shear stress in the cell membrane in gradient and uniform MFs, accordingly. Both stresses depend on the MF direction and magnitude. For the certain directions of an uniform MFs the shear stress is zero: (*θ* = 0, *θ* = *π*), (γ = 0 or γ = *π*).

In a gradient MF, shear stress (Eq. 5) can close/open the membrane ion channels. In such a case, for high MF gradients, the effect of “magnetic saturation of channels” is expected: a high enough magnetic shear stress keeps ion channels in the fully opened state [82].

Let us introduce the upper limit of the WSS which keeps the channels in the open state. Strictly speaking, as dictates Eq. 2, a channel is fully open when the energy associated with shear stress, *W* goes to infinity. In this case, we can define the open state probability, for example as *f*_*0*_=0.9. Thus, by this definition, 90 % of channels are open at the certain value of shear stress, which we call as stress of the channel saturation, *P*_s_. See Supplementary Material (Fig. S4) for estimation of the Ca^2+^ channel saturation reached at *P*_s_≈ 5 Pa in endothelial cells.

Now, we introduce two important dimensionless parameters that control intracellular calcium dynamics through magnetic shear stress, which drives mechanosensitive Ca^2+^ ion channels. The first parameter is 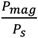 where the expression *P*_*mag*_ is the amplitude of magnetic WSS (for definition see Section 4, Methods), and *P*_*s*_ is the WSS corresponding to saturation of the current through the mechanosensitive Ca^2+^ ion channel assuming the probability of channel opening *f*_*o*_ = 0.9 in Plank’s model. The parameter *m* is used for both cases: WSS induced by a gradient MF (Eq. 5) and for WSS induced by a uniform MF (Eq. 6).

The second parameter is 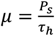, where τh is WSS due to blood flow in a vessel, e.g., τ_*h*_=1Pa for arteryand τ_*h*_= 0.1 Pa [79] for capillary. The *μ*-parameter is independent of a MF and serves as a material parameter: μ≈5 for artery and μ≈50 for capillary.

Both material and magnetic parameters (*μ* and *m*) are introduced in the calcium dynamics governing equations (S8-S11). These equations were numerically solved using Python programming language, numpy and scipy packages. The results on Ca^2+^ dynamics are presented in the (*m, ν*)-coordinates for various values of the μ-parameter, which is associated with different endothelial cells, and the frequency of an oscillating MF, ν. The results of calculations and discussions are provided for the set of parameters from Plank’s model [28], which describes the gating of Ca^2+^ ion channels and intracellular calcium dynamics in endothelial cells (Table S2).

### Endothelial Ca^2+^ dynamics

Vascular endothelial cells, which form the inner lining of the blood vessel wall and are directly exposed to blood flow and corresponding shear stress, serve crucial homeostatic functions in response to various chemical and mechanical stimuli [82,83].

To reveal the effects of low-frequency MF on the concentration of free Ca^2+^ in the cytosol, we numerically solve Eqs. S8-S11 (in the Supplementary Materials) with respect to Eqs. 4-6. Note, equations S8-S11 represent the dimensionless forms of equations 7-10, as provided in the Material and Methods section. The results of calculations are mapped onto the (*m,ν*)-diagrams (Fig. 3). In Fig. 3, we plot the following characteristics of calcium dynamics for different MF amplitudes (related to the *m*-parameter) and MF frequencies (ν): the maximum amplitude of spike of free cytosolic calcium concentration 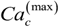, the time averaged amplitude of spike of free cytosolic calcium concentration 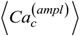, the time averaged concentration of free cytosolic calcium ⟨*Ca*_*c*_⟩, and the time averaged frequency of quasiperiodic spikes of free cytosolic calcium concentration 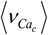 (Fig 3. E and F). It is worth noting that the selected range of MF frequencies, ranging from 6 mHz to 17 mHz, corresponds to the optimal specific frequencies of decoding enzymes, including 1) MAPK, 2) NF-κB, 3) NFAT, and 4) glycogen phosphorylase kinase [5]. It is important to note that the frequency range (approximately 10 mHz) of the decoding enzymes is determined by molecular motor dynamics, in which the typical binding rates fall within the range (1-5) 10^-2^ s^-1^ [84]. The upper limit of the calcium signaling frequency is related to the time, Δt=1/ν, which corresponds to the mean time of calcium waves propagations in a cell. The upper limit can be estimated as ν_max_=u/L ≈ 20 μm s^-1^/20 μm ≈ 1 Hz for L= 20 μm and the velocity of calcium waves, *u*= 20 μm s^-1^ [85]. In the next sections we will continue discussing the results presented in Fig. 3.

**Fig. 3.**
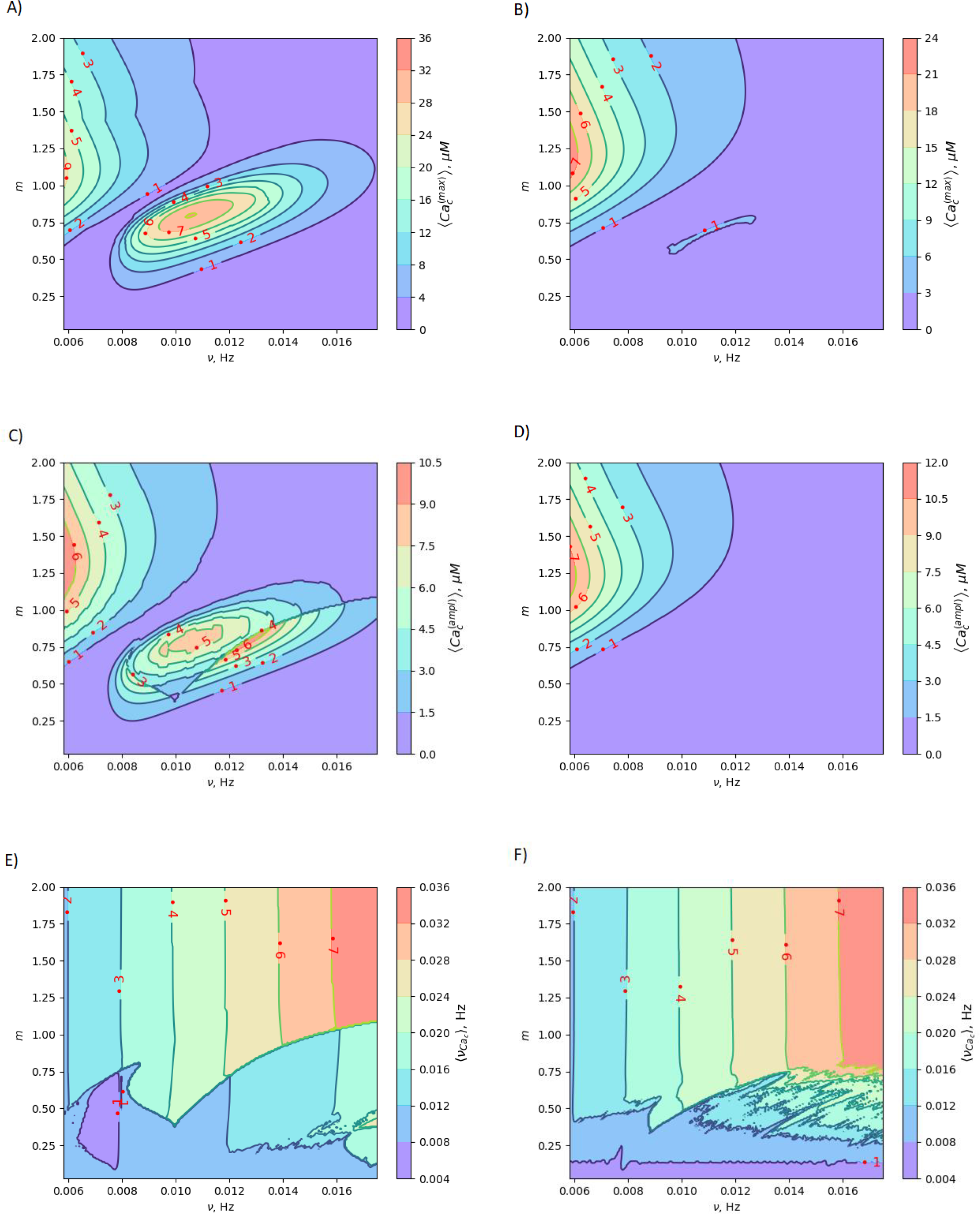
Mapping MF effects on dynamics of intracellular free Ca^2+^ concentration in the vascular endothelium in low-frequency MF. Here, *ν* is the MF frequency; 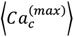 is maximum amplitude of the spike of free cytosolic Ca^2+^ concentration for artery; B) 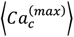is maximum amplitude of the spike of free cytosolic Ca^2+^ concentration for capillary; C) 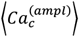is time averaged amplitude of the spike of free cytosolic Ca^2+^ concentration for artery; D) 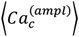is time averaged amplitude of the spike of free cytosolic Ca^2+^ concentration for capillary; E) 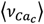 is time averaged quasi-frequency of oscillations of free cytosolic Ca^2+^ concentration for artery; F) 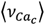 is time averaged quasi-frequency of oscillations of free cytosolic Ca^2+^ concentration for capillary. The oscillatory regime of intracellular calcium dynamics is considered before application of MF (the set of parameters used in the calculations for oscillatory regime is also listed in the last 5^th^ column of Table S2 in the Supplementary Materials).

### Sensing frequencies of decoding proteins

In human aortic endothelial cells, the frequency of agonist-induced Ca^2+^ oscillations is positively correlated with the NF-κB activity in the range of 1.8–5.3 mHz, with a duration of approximately 160 s (duty cycle 0.29%– 0.85%) [5]. Using another experimental setup with Ca^2+^ clamping in combination with agonist stimulation, vascular endothelial cells were shown to regulate VCAM1 expression with a Ca^2+^ frequency in the range of (1.7–11.7) mHz, with a duration of approximately 29 s (duty cycle 0.049%–0.34%) [5]. In human cerebral endothelial cells treated with sarco/endoplasmic reticulum Ca^2+^-ATPase and inositol 1,4,5-trisphosphate receptor (InsP3R) inhibitors to modulate the Ca^2+^ frequency, the NF-κB activity is positively correlated with frequencies in the range of (0–5.2) mHz (duration 100 s) [5]. Histamine was also used in human aortic endothelial cells showing that the NF-κB activity is positively correlated with Ca^2+^ frequency in the range of (2–4) mHz (duration 60 s) [5]. There are the following decoding proteins of Ca^2+^ oscillations in endothelial cells: NFAT, NF-κB, MAPK and glycogen phosphorylase kinase [5]. The sensing frequencies of these proteins are given in [5] and listed below. MAPK (mitogenactivated protein kinase). Ca^2+^ frequencies in the range of (1.7–17) mHz with a duration of approximately 50 s (duty cycle 0.085%–0.85%). NF-κB (nuclear factor kappa-light-chain-enhancer of activated B cell) the Ca^2+^ frequency in the domain of (0.56–10) mHz, with a duration of approximately 100 s (duty cycle 0.028%–0.5%) [5]. Notably, the minimum frequency is 4.5 times lower than that for NFAT [5]. The main difference between NFAT and NF-κB is the higher sensitivity of NF-κB, where frequencies as low as 0.56 mHz suffice [5]. NF-κB seems to be tuned toward higher duty cycles (0.8%–0.9%) than NFAT. The optimal frequency is around 10 mHz [5]. NFAT (nuclear factor of activated T-cells) is activated by the phosphatase calcineurin, frequencies in the range of (2.5–10) mHz, with a duration of 50 s (duty cycle 0.125%–0.5%). The maximum NFAT activity is present at the Ca^2+^ frequency of 16.7 mHz (duty cycle 0.33%) with decreasing activity down to 2 mHz and up to 33 mHz, with a duration of 20 s (duty cycle 0.04%–0.66%). The optimal frequency is approximately 20 mHz and duty cycle around 0.2–0.3. GP (Glycogen phosphorylase) is activated by a kinase that has a calmodulin-like Ca^2+^ sensitive domain, frequency interval of (1.7–170) mHz, whereas the enzyme shows constant activity at higher and lower frequencies [5].

### Superharmonic resonance and shift of the Ca^2+^ spiking frequency in MFs

The (*m, ν*) - diagrams (Fig. 3) demonstrate that the MF influence on intracellular Ca dynamics in both oscillatory and non-oscillatory regimes (Fig. S3 A, Fig. S3 B). The appearance of either the oscillatory or non-oscillatory regime is determined by the actual values of both parameters: *m* and μ, and parameters of the Plank’s model not dependent on MF (Table S2). The Plank’s model posits both oscillatory and non-oscillatory regimes in intracellular dynamics [28]. Different parameters in the Plank’s model are applicable to the oscillatory regime (presented in the last 5th column of Table S2 in the Supplementary Material) and the non-oscillatory regimes of intracellular dynamics (presented in the 4th column of Table S2 in the Supplementary Material). Both oscillatory and non-oscillatory regimes can occur within the same ranges of the *m* and *v* parameters, as the switch between these regimes is exclusively controlled by the Plank’s model parameters without a magnetic field. However, the resulting influence of a magnetic field on intracellular calcium dynamics differs for these two regimes. In the non-oscillatory regimes (in Fig. S3 in the Supplementary Material), the average Ca^2+^ concentration and its standard deviation from the average value doesn’t depend on the MF frequency, ν.

In the oscillatory regimes, the average Ca^2+^ concentration, the amplitude and frequency of Ca oscillation depend on the MF frequency in a resonant-like manner (Fig. 3, Fig. 4 (A, B, C, D, E, F)). In Fig. 3, the diagrams show the frequency ranges for which the magnetic field effects on Ca^2+^ concentration are more pronounced. The significant impact on Ca dynamics from oscillating uniform and gradient magnetic fields (with the *m*-parameter ranging from 0.25 to 2.0) occurs at low frequencies comparable to the frequency of Ca concentration self-oscillations in the absence of a magnetic field. This finding qualitatively aligns with experimental data [86], where the impact of magnetic fields on the spectral power of the cytosolic Ca^2+^ oscillations was observed at lower frequencies, specifically within the (0–10) mHz subinterval of the Ca^2+^ oscillation spectrum. However, for the high frequency ranges (Fig. 3), the change Ca concentrations caused by the MFs (Fig. 5 (A, B, C, D, E, F)) are not significant, which also in a good agreement with experiments [86].

**Fig. 4.**
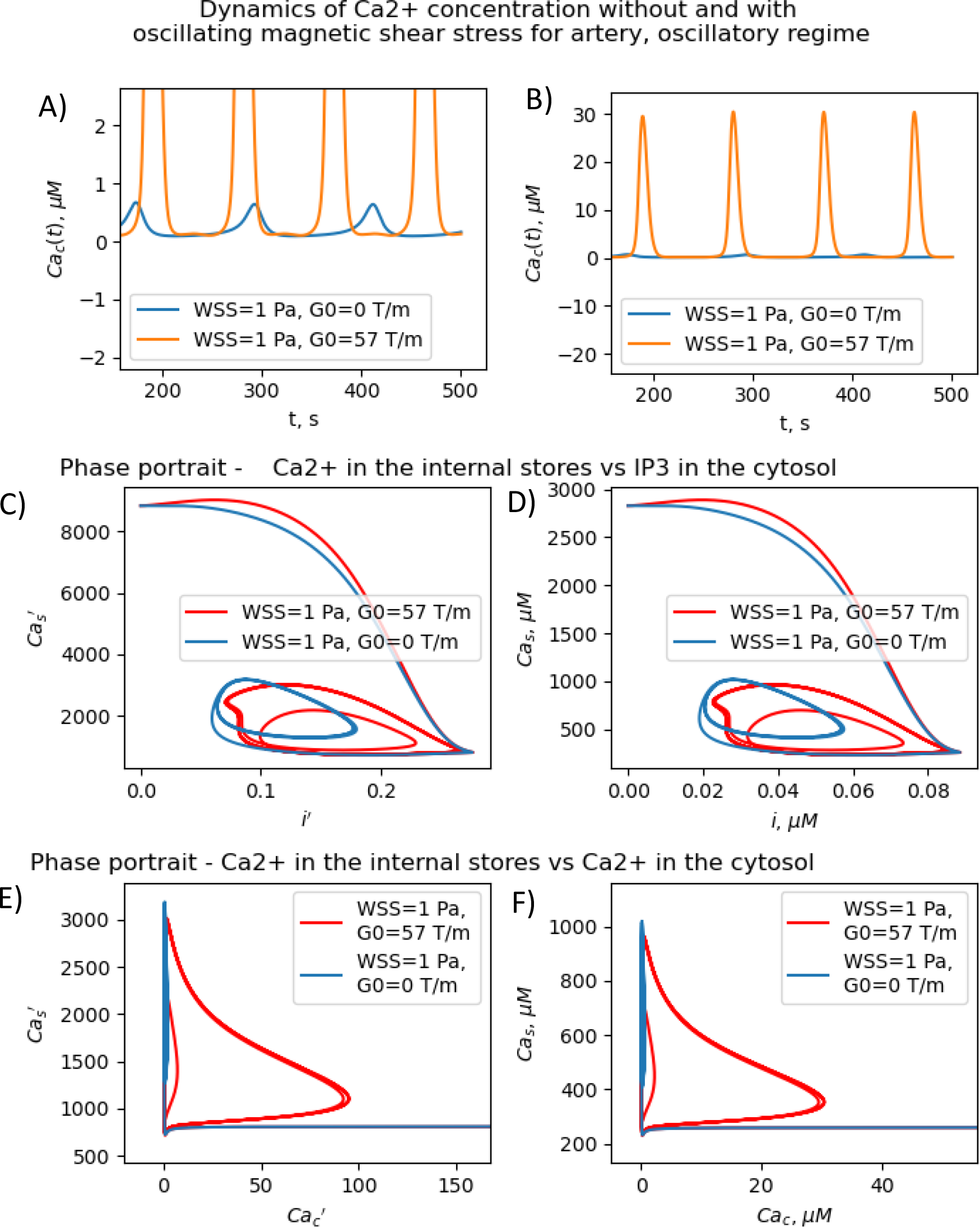
Resonant-like dynamics of concentration of intracellular Ca^2+^ for artery, *m* = 0.77, *ν* = 0.011 Hz in a gradient MF. (A)Temporary dependencies of the dimensionless concentration of free calcium in the cytosol with and without the gradient MF. The gradient MF with frequency ν = 0.011 Hz significantly increases the amplitude of calcium oscillations.(B)Temporary dependencies of the dimensionless concentration of free calcium in the cytosol with and without the gradient MF. The gradient MF with frequency = 0.011 Hz significantly increases the amplitude of calcium oscillations.(C)Dependence of the dimensionless concentration of calcium in the internal store on the dimensionless concentration of IP3 with and without the gradient MF (*m* = 0.77, *ν* = 0.011 Hz in artery). D) Dependence of the dimensionless concentration of calcium in the internal store on the dimensionless concentration of IP3 with and without the gradient MF (*m* = 0.77, *ν* = 0.011 Hz in artery). E) Dependence of the dimensionless concentration of calcium in the internal store on the dimensionless concentration of IP3 with and without the gradient MF (*m* = 0.77, *ν* = 0.011 Hz in artery).(F)Dependence of the dimensionless concentration of calcium in the internal store on the dimensionless concentration of IP3 with and without the gradient MF (*m* = 0.77, *ν* = 0.011 Hz in artery).

**Fig. 5.**
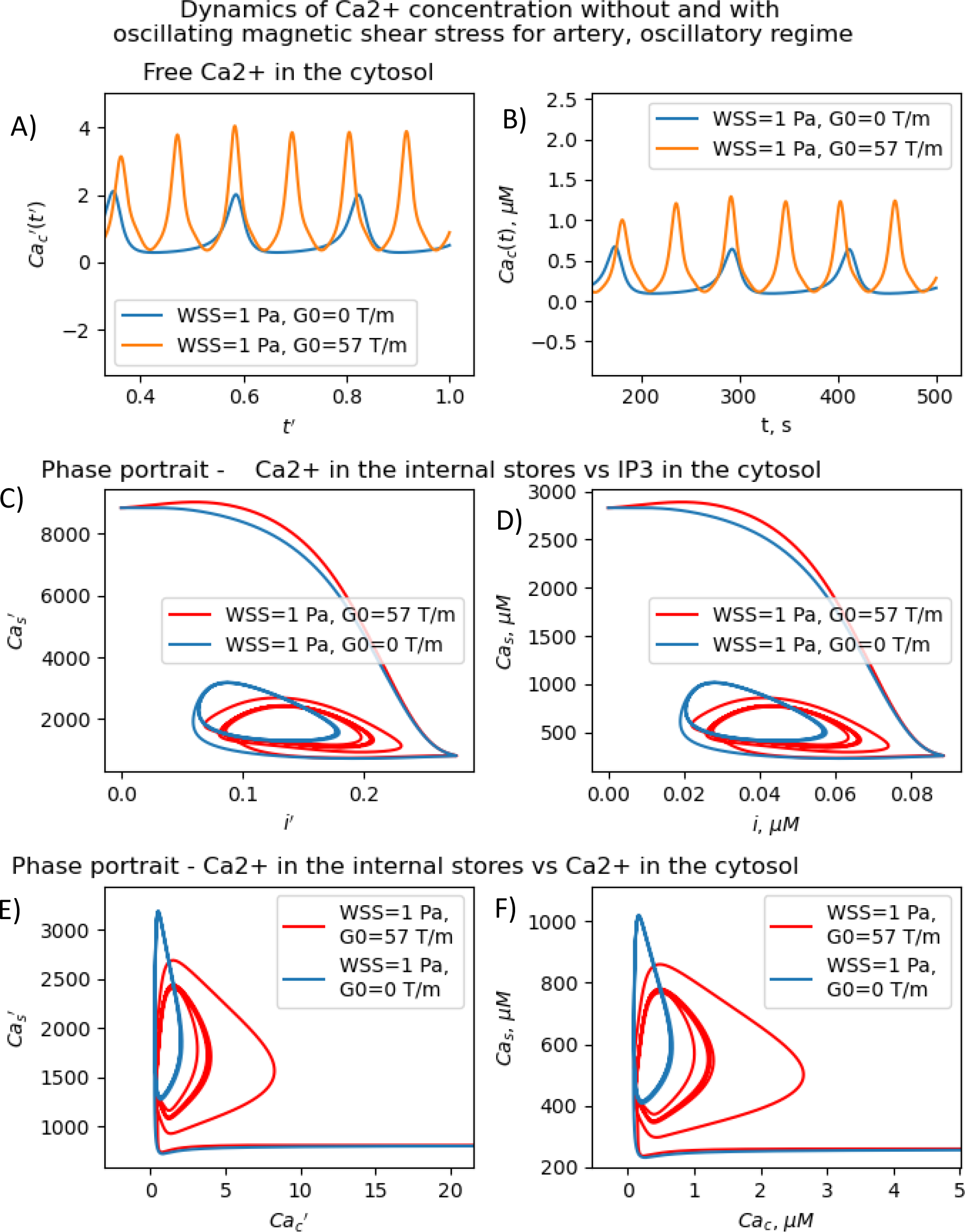
Non-resonant dynamics of concentration of intracellular Ca^2+^ for artery in gradient MF (*m* = 0.77, *ν* = 0.018 Hz).(A)Temporary dependence of the concentration of free calcium in the cytosol. The magnetic field increases the amplitude of calcium oscillations and doubles their frequency. B) Temporary dependence of the dimensionless concentration of free calcium in the cytosol. The magnetic field increases the amplitude of calcium oscillations and doubles their frequency. C) Dependence of the dimensionless concentration of calcium in the internal store on the dimensionless concentration of IP3 with and without the gradient MF (*m* = 0.77, *ν* = 0.018 Hz in artery). D) Dependence of the dimensionless concentration of calcium in the internal store on the dimensionless concentration of IP3 with and without the gradient MF (*m* = 0.77, *ν* = 0.018 Hz in artery). E) Dependence of the dimensionless concentration of calcium in the internal store on the dimensionless concentration of IP3 with and without the gradient MF (*m* = 0.77, *ν* = 0.018 Hz in artery). F) Dependence of the dimensionless concentration of calcium in the internal store on the dimensionless concentration of IP3 with and without the gradient MF (*m* = 0.77, *ν* = 0.018 Hz in artery).

In Fig. 3, the diagrams display characteristic traits associated with both the *foldover effect* and *superharmonic resonance*. In general, they represent distinct features of nonlinear oscillators that do not oscillate sinusoidally. In such a system, oscillation is not a single sinusoidal wave, but rather a combination of sinusoidal waves with frequencies that are whole number multiples of the fundamental frequency. In our case, the first resonant frequency, *ν* ≈ 0,009 Hz in Fig. 3 A, C corresponds to the frequency of the first maximum in amplitude spectrum of the temporal dependence of the free Ca^2+^ concentration for an artery without MF as obtained with a discrete Fourier transform after the subtraction of the average Ca^2+^ concentration (see Supplemental Material at [URL will be inserted by publisher] for *.zip file containing the python project *MagniCa*). The second resonant frequency, *ν* ≈ 0,013 Hz (Fig. 3 A, C) corresponds to the frequency of the second maximum in the amplitude spectrum of the temporal dependence of the free Ca^2+^ concentration for the artery without MF based on a discrete Fourier transform after the subtraction of the average Ca^2+^ concentration.

In Figs. 3, in (A, C, E) diagrams calculated for arteria, for the intermediate MF frequencies (10-12 mHz) and the *m*-parameter lying in the range 0.75 -1.0, one can see the domain in which the amplitude of the cytosolic calcium oscillation reaches its maximum. For capillaries (Figs. 3, (B, D, F)), in the same frequency and *m* domain, the cytosolic calcium concentration is only slightly increased under the MF influence. The average frequencies of Ca^2+^ ion channel gating as a function of *m* and ν (which is MF frequency) are shown in Fig. 3 E and Fig. 4 F, for arteries and capillaries, accordingly. There is the threshold represented by the *m(ν)-*curve, upon reaching which the oscillation frequency of cytosolic Ca^2+^ sharply doubles. Physically, the doubling of Ca^2+^ oscillation frequency is associated with the saturation of ion currents through Ca channels on reaching *m*=1 (Fig. 3 E, Fig. 3 F), which, in turn, is a direct consequence of the effect of Ca^2+^ ion current saturation under high WSS (a high enough magnetic shear stress (Eq. 5) keeps ion channels in the fully opened state [82]. In the other words, if *m* approaches unity the magnetic WSS value achieves the saturation stress, *P*_*s*_ = 5 *Pa* at which the Ca^2+^ ion channels are open with probability 0.9. For example, the ion current saturation takes place in MFs with the gradient *G*_0_ ≈ 80 T/m and BMNs with *r* = 100 nm, *M*_*s*_ = 510 · 10^3^ A/m. In both cases, for arteria (Fig. 3 E) and capillaries (Fig. 3 F), after the frequency doubling, the further increase of a MF or its gradient doesn’t lead to an increase of the rate of cytosolic Ca^2+^ oscillations and doesn’t affect intracellular Ca dynamics (Fig. 3, E and F).

The MF influence on the gating of mechanosensitive Ca^2+^ ion channels and intracellular Ca dynamics is low if the first magnetic parameter *m* ≪ 1. This conclusion is valid only for non-oscillatory dynamics of Ca^2+^ concentration and for oscillatory dynamics of Ca^2+^ concentration in the case when a MF frequency is high in comparison with the resonant one. For example, *m* = 0.014 where *G*_0_ =1T/m, *r* = 100 nm, *M*_*s*_ = 510 · 10^3^ A/m, *P*_*s*_ = 5 *Pa*. In the last case, the MF influence on the probability of opening of mechanosensitive Ca^2+^ ion channels is about 1-2% and the MF influence on the rate of intracellular Ca flux inside a cell is about 2% in comparison with the conditions without MF.

In contrast, for oscillatory Ca^2+^ concentration dynamics, notable effects on intracellular Ca dynamics are observed even with small *m-*parameters. These effects become particularly pronounced when exposed to low gradient MFs and low uniform MFs, especially when the frequency is lower than the self-oscillation frequency of Ca^2+^ concentration. Indeed, the diagrams (Fig. 3) show that the MF changes the concentration of intracellular free calcium from resting value (non-stimulating around 0.1 µM=100 nmol/L) to the level of several µM activating signaling transduction. That is why an MF of appropriate strength and frequency induces additional WSS and renders an influence on calcium dependent processes.

The results presented in Figs. 3-5 show that the stable concentration of Ca^2+^ level is established not immediately but after some delay after application of magnetic field (transition processes) in the case of non-oscillatory regime of Ca^2+^ dynamics. A delay (transition processes) also appears in the oscillatory regime of Ca^2+^ dynamics (Fig. 4, Fig. 5). This means that it takes some time for self-oscillations to occur under the influence of a magnetic field. The transition process is slow, the delay time is about hundreds of seconds. This prediction of the model is in an agreement with the observation of slow relaxation of brain activity under the influence of a magnetic field (with magnetic field flux density close to the geomagnetic field) [87], namely, alpha-power began to drop from pre-stimulus baseline levels as early as 100 ms after magnetic stimulation, decreasing by as much as 50% over several hundred milliseconds, then recovering to baseline by 1 s post-stimulus [87].

The present analysis sheds new light on the longstanding issue of the impact of magnetic field on membrane ion channels. Actually, we show that, by applying low oscillating gradient magnetic fields to endothelial cells carrying chains of biogenic or artificial magnetic nanoparticles, the resonance phenomenon and doubling frequency of calcium oscillations are found (Figs. 4-5). A superharmonic resonance, accompanied by a sharp increase in the amplitude of calcium oscillations, takes place when the MF frequency is close to the self-frequency of free calcium oscillations in the absence of a magnetic field. Here, the magnetic field-amplified calcium concentration and the magnetic field-induced frequency shift of calcium oscillations result in the modulation of both amplitude and frequency of the calcium waves. The frequency modulation of the calcium waves propagating in the endothelium is of special interest.

It is known that, in endothelial cells, Ca waves can propagate within arterial wall [88]. Since Ca waves are often initiated by intracellular release of calcium from stores such as the endoplasmic reticulum, they can be triggered by a MF in arteria with the above-mentioned values of the *m* and μ parameters. This provides an opportunity for frequency modulation of calcium waves using a magnetic field, which could enable the control of various cellular processes, including cell proliferation, differentiation, and migration. Fig. 6 schematically demonstrates how to alter cell fate by magnetic switching of the frequency bands of calcium waves. Detailed information regarding the MF frequency bands, decoding of proteins, and metabolic pathways is provided in Table 3.

**Fig. 6.**
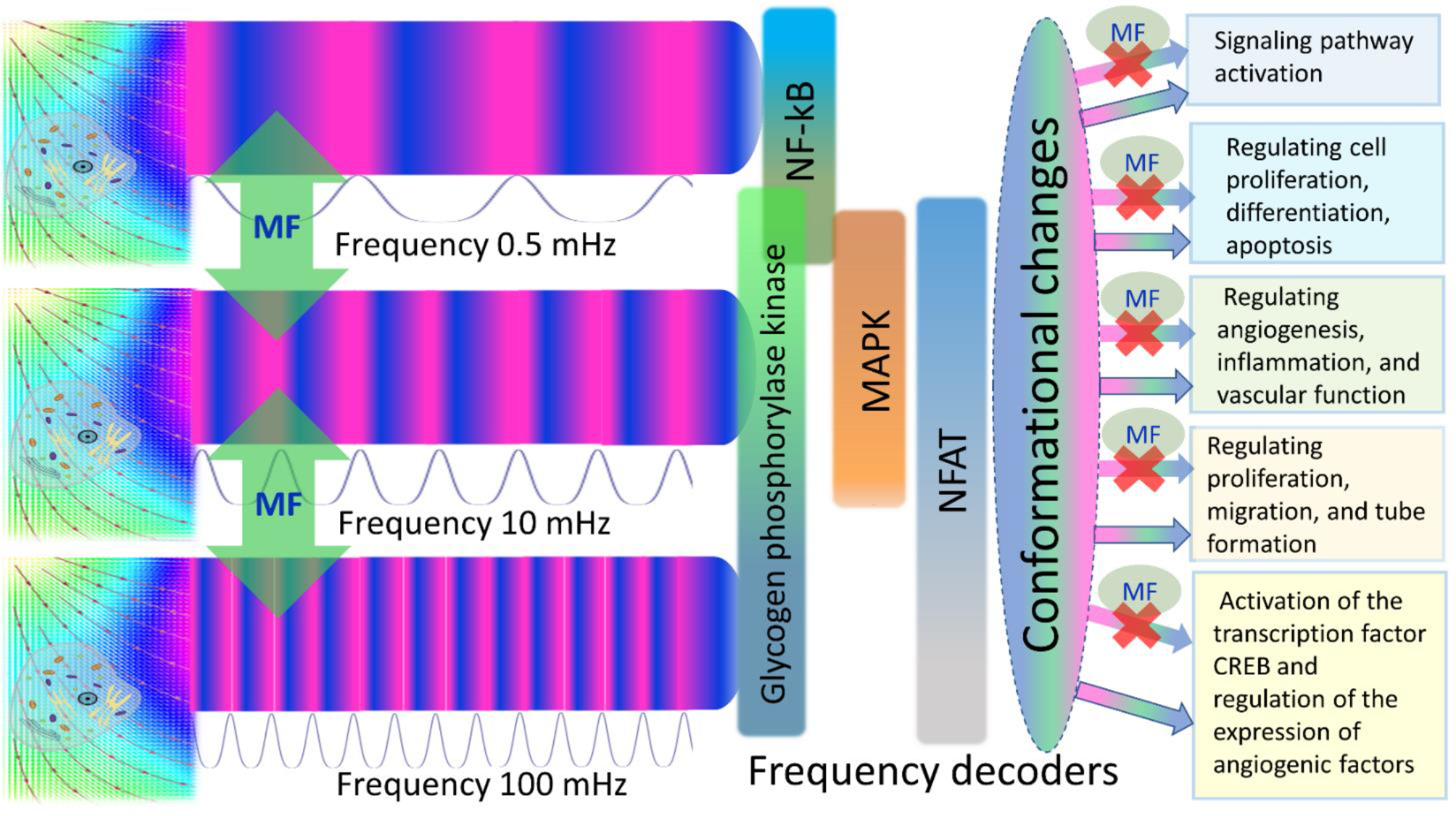
Frequency modulation of calcium waves and magnetic switching of metabolic pathways in endothelial cells is depicted with four frequency decoders shown: NF-kB, MAPK, NFAT, and Glycogen phosphorylase kinase (GPK) [5]. A magnetic field (green vertical arrows and spots marked with MF) switches the frequency bands of calcium waves, thereby changing/closing enzyme activity and metabolic pathways.

## Discussion

### Regulation of cell fate with magnetic fields

We suggested an approach for modulating the calcium signaling frequency using uniform and gradient magnetic fields. Our theoretical model is based on the previous experimentally verified models of calcium dynamics in endothelial cells [28,29]. The key points of our approach are the following. An externally applied MF exerts forces on biogenic and non-biogenic magnetic nanoparticles which are always present in organisms (Fig. 1). The nanoparticles transmit magnetic forces to the cell membrane, causing shear stress in cell walls, which, in turn, changes the activity of calcium ion channels and modifies the intracellular calcium pattern. The application of a time-varying magnetic field enables both frequency modulation of calcium signaling and switching between calcium decoders. From this perspective, it is believed that certain human diseases resulting from abnormal intracellular calcium levels and disruptions in calcium signaling may be treated with time-varying magnetic fields. The specific frequency and amplitude of these fields can be determined by two parameters: *m* and μ, which operating values are depicted in Figs. 3-5. Finally, the possibility of the magnetic switching between the calcium decoders (Fig. 6) opens the door for the modulation of biochemical pathways and the fate of human cells in the absence of chemical or biological agents.

### Treatment of diseases with alternating magnetic fields

Since elevated calcium level stimulates endothelial proliferation [89] and promotes the growth of new vessels (angiogenesis) [89], a MF can serve as noninvasive tool for calcium level setting. In this view, static and alternating MFs can be used for treating injuries, trauma, strokes; enhancing wound healing (if magnetic shear stress is applied along the hydrodynamic one) or for inhibiting endothelial proliferation for anticancer treatment (if magnetic shear stress is applied opposite to the hydrodynamic one. In cases when elevated calcium leads to rise of blood pressure [90], the magnetic field induced shear stress, which is parallel to hydrodynamic (blood flow), can be used for treating hypotension and magnetic shear antiparallel to blood flow, to treat hypertension. So far as an elevated calcium concentration results in increase of endothelial permeability [91], in particular, supporting transendothelial migration of immune cells [92], a MF can be utilized for immunomodulation. For example, a magnetic field generating magnetic shear stress, which opposes the hydrodynamic force, can be used to decrease endothelial permeability for immune cells. Consequently, this can inhibit inflammation and aid in the development of immunosuppressive drugs for organ transplantations, as well as the treatment of autoimmune diseases, among other applications. Moreover, calcium signaling in endothelial cells induces neuromodulation and impacts blood flow control [93]. Hence, the application of a MF shows promise for controlling neurovascular coupling and treatinFg vascular dysfunctions associated with aging and neurodegenerative disorders. Future research in the direction of magnetic field induced wall shear stress and gating of calcium ion channels in endothelial cells opens perspective to treat cardiovascular diseases (including atherosclerosis) and cancer.

MF-assisted Ca^2+^ ion channel gating can also be utilized to induce magnetic field-assisted synchronization of calcium oscillations in all cells within a target region (organ or tissue). This approach can be used to treat diseases in which defects in this coordinating mechanism contribute to the pathology, for example, Type 2 diabetes [94].

The results of this research contribute to the targeting of ion channels, which represent the second-largest class of membrane proteins in the market after G Protein-Coupled Receptors (GPCRs) [25]. To achieve that, we propose the administration of chains of artificial and/or biogenic magnetic nanoparticles to a specific lesion site, followed by the application of appropriate magnetic shear stress as a therapeutic method for targeting the mechanosensitive Ca^2+^ ion channels in endothelial cells.

Summarizing, our findings might pave the way for the use of oscillating and pulsed magnetic fields to improve functions of endothelial cells. The suggested model and obtained results are of great importance for further developing novel noninvasive and nondestructive physical approaches in cell therapy and medicine.

### Possible hidden mechanisms of biomagnetic effects

Thousand papers dealing with study of biomagnetic effects of low and moderate MFs do not pay attention to a possible presence of biogenic and residual artificial magnetic nanoparticles in organisms, in particular, in endothelial cells [45,95–97]. Nevertheless, as above discussed such nanomagnets are the active targets of static and especially low-frequency oscillating and rotating MFs. In this view, the above suggested physical mechanism and model of the magnetic field impacts on the calcium patterns and signaling in endothelial cells can play a hitherto unexpected role in creating physiological responses of organisms to externally applied MFs. The range of individual responses may be partially attributed to diversity in nanoparticles number and their localization rather than to other underlying processing.

### Prospects for experiments

Future experiments should examine how static and alternating MFs interact with other ion channels on cell membranes bound to biogenic and artificial residual magnetic nanoparticles to determine mechanical stresses and mechanisms of changing cell machinery. Future studies should also examine differences in responsiveness of different cell types to magnetic fields and signal processing. It is also highly desirable to develop theoretical models to investigate the effects of magnetic fields on calcium dynamics in other cell types, especially MF impact on calcium signaling in brain neurons. We hope that our study provides a roadmap for future studies aiming to replicate and extend research into magnetobiology.

## Materials and methods

### Model

Plank’s model [28] of intracellular calcium dynamics was generalized, taking into account the wall shear stress (WSS) induced by a gradient or uniform magnetic field, as well as a static or oscillating magnetic field. This was achieved by considering the magnetic field’s influence on biogenic and/or artificial magnetic nanoparticles embedded in the cell membrane of endothelial cells. The WSS induced by a gradient or uniform magnetic field is calculated according to formulas (5) and (6). The MF-induced WSS changes the probability of Ca^2+^ mechanosensitive channel opening and the rate of Ca^2+^ flux inside the cell according to formulas (2) and (4), respectively.

### The main equations, notions and terms

The Plank’s model of Ca^2+^ dynamics considers 4 dynamic variables: concentrations of IP3 in the cytosol, free Ca^2+^ in the cytosol, buffered Ca^2+^ in the cytosol, Ca^2+^ in the internal stores. The numerical values of the dynamic variables of the model of Ca^2+^ dynamics are collected in Table 1 and Table S1.

**Table 1.** Dynamic variables in the modified Plank’s model for calcium dynamics taking on account WSS induced by magnetic field.

| Dynamic variable | Dimensional notation |
| --- | --- |
| Concentrations of IP3 in the cytosol | $i$ |
| Free $\text{Ca}^{2+}$ in the cytosol | $Ca_c, \mu M$ |
| Buffered $\text{Ca}^{2+}$ in the cytosol | $Ca_b, \mu M$ |
| $\text{Ca}^{2+}$ in the internal stores | $Ca_s, \mu M$ |
| Sum of free $\text{Ca}^{2+}$ in the cytosol and buffered $\text{Ca}^{2+}$ in the cytosol | $Ca_t = Ca_c + Ca_b, \mu M$ |

The governing equations for modelling calcium dynamics [28] for the dynamic variables represented in Table 1:

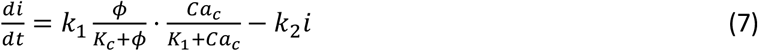

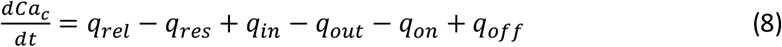

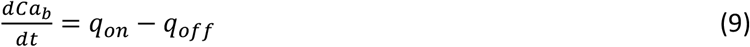

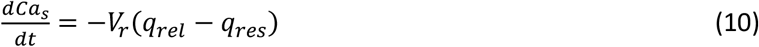

Note that in Eq. 8, there is one parameter, *q*_*in*_, which is dependent on the magnetic field through the wall shear stress. The variables, rates and parameters in the right parts of (7)-(10) are represented in Table 1, Table 2,Table S1, and Table S2. See Supplemental Material at [URL will be inserted by publisher] for the transformation of the system of equations (7)-(10) to the dimensionless form.

**Table 2.**
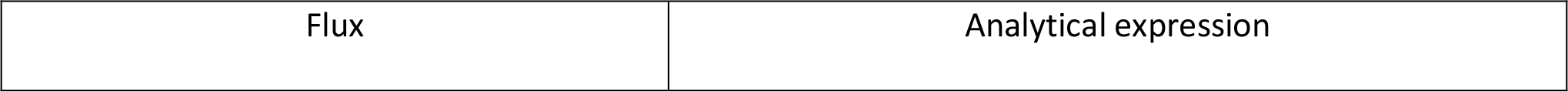
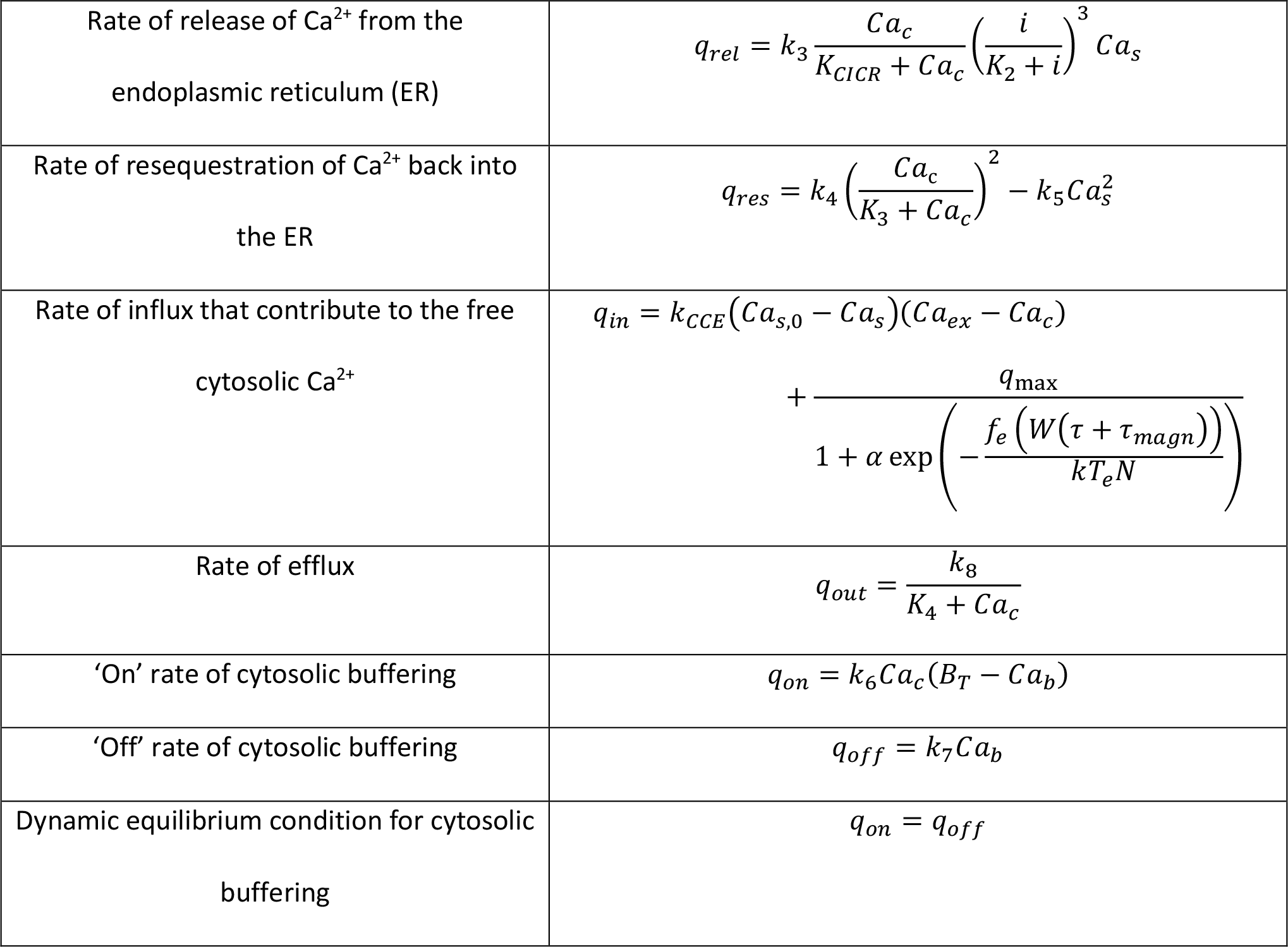
Fluxes in the modified Plank’s model for calcium dynamics taking on account WSS induced by magnetic field. The parameters in the second column are represented in Table S2.

**Table 3.** Magnetic field impacts on decoding free intracellular calcium signals in endothelial cells.

| Decoding protein<br>• Metabolic pathway / Description<br>• Diseases related to dysregulation of the pathway, drug targets | Frequency range for effective decoding /<br>Duty cycle range for effective decoding<br>• optimal frequency / optimal duty cycle<br>• Magnetic field effects |
| --- | --- |
| MAPK (mitogen-activated protein kinase), mitogen-activated protein kinase 4 (Map4k4)<br>• MAPK signaling pathway / The MAPK/ERK pathway (also known as the Ras-Raf-MEK-ERK pathway) is a chain of proteins in the cell that communicates a signal from a receptor on the surface of the cell to the DNA in the nucleus of the cell. The signal starts when a signaling molecule binds to the receptor on the cell surface and ends when the DNA in the nucleus expresses a protein and produces some change in the cell, such as cell division. The pathway includes many proteins, such as mitogen-activated protein kinases (MAPKs), originally called extracellular signal-regulated kinases (ERKs), which communicate by adding phosphate groups to a neighboring protein (phosphorylating it), thereby acting as an "on" or "off" switch. MAPK or MAP kinase is a type | • 1.7–17 mHz /<br>• 0.085–0.85<br>• |
| of protein kinase that is specific to the amino acids serine and threonine (i.e., a serine/threonine-specific protein kinase) [101]. MAPKs are involved in directing cellular responses to a diverse array of stimuli, such as mitogens, osmotic stress, heat shock and proinflammatory cytokines [101]. They regulate cell functions including proliferation, gene expression, differentiation, mitosis, cell survival, and apoptosis. MAP kinases are found in eukaryotes only, but they are fairly diverse and encountered in all animals, fungi and plants, and even in an array of unicellular eukaryotes [101].<br>• When one of the proteins in the pathway is mutated, it can become stuck in the "on" or "off" position, a necessary step in the **development of many cancers**. Components of the MAPK/ERK pathway were first discovered in cancer cells, and drugs that reverse the "on" or "off" switch are being investigated as cancer treatments. Constitutive loss of endothelial cell Map4k4 in mice causes postnatal lethality due to |  |
| **chylothorax**, suggesting that Map4k4 is required for normal lymphatic vascular function [102]. Mice constitutively lacking EC Map4k4 displayed **dilated lymphatic capillaries, insufficient lymphatic valves, and impaired lymphatic flow** [102]; furthermore, primary endothelial cells derived from these animals displayed enhanced proliferation compared with controls [102]. |  |
| NF-κB (NFKB in KEGG - nuclear factor NF-kappa-B; nuclear factor kappa-light-chain-enhancer of activated B cells)<br>The Nuclear Factor NF-κB Pathway / The nuclear factor NF-κB pathway has long been considered a prototypical proinflammatory signaling pathway, largely based on the role of NF-κB in the expression of proinflammatory genes including cytokines, chemokines, and adhesion molecules. NF-κB is a protein complex that controls transcription of DNA, cytokine production and cell survival [103–105]. NF-κB is found in almost all animal cell types and is involved in cellular responses to stimuli such as stress, cytokines, free radicals, | 1) 0.56–10 mHz / 0.028–0.5<br>2) 0.56 mHz / 0.8–0.9<br>10 mHz |
| heavy metals, ultraviolet irradiation and bacterial or viral antigens. NF-κB plays a key role in regulating the immune response to infection. NF-κB target genes could be classified in different functional groups [103]. Those with a primarily inflammatory function were predominant [103]. NF-κB controls the global pro-inflammatory response in endothelial cells [103]. Genetic evidence in mice has revealed complex roles for the NF-κB in inflammation that suggest both pro- and anti-inflammatory roles for this pathway. NF-κB has also been implicated in processes of synaptic plasticity and memory.<br>• Incorrect regulation of NF-κB has been linked to **cancer, inflammatory and autoimmune diseases, septic shock, viral infection, and improper immune development**. NF-κB has long been considered as a **target for new anti-inflammatory drugs** [105]; however, these recent studies suggest this pathway may prove a difficult **target in the treatment of chronic disease** [103,105]. Deregulated NF-κB activation contributes to the pathogenic |  |
| processes of various inflammatory diseases [106]. |  |
| NFAT (Nuclear Factor of Activated T-Cells)<br>Calcium–Calcineurin–NFAT Signaling Pathway; $\text{Ca}^{2+}$/nuclear factor of activated T-cells ($\text{Ca}^{2+}$/NFAT) signaling pathway / NFAT signaling plays a key role in angiogenic cell behaviors. NFAT is a family of transcription factors shown to be important in immune response [107]. One or more members of the NFAT family is expressed in most cells of the immune system. NFAT is also involved in the development of cardiac, skeletal muscle, and nervous systems [107]. NFAT was first discovered as an activator for the transcription of IL-2 in T cells (as a regulator of T cell immune response) but has since been found to play an important role in regulating many more body systems [107]. NFAT-mediated gene upregulations lead to well-known T cell differentiations and activation, but most recently, many reports also showed the NFAT activity correlates with innate immune cell activation, bone metabolism, angiogenesis, and tissue | 1) 2.5–10 mHz / 0.125–0.5<br>2) 2 – 33 / 0.04–0.66<br>3) 16.7 mHz / 0.33<br>20 mHz / 0.2–0.3 |
| regenerations [108]. In the vascular system NFAT (including the isoforms c1 and c3) contributes to cell growth, remodeling of smooth muscle cells, controls vascular development and angiogenesis and is activated in response to inflammatory processes and high intravascular pressure [109]. In the endothelium, NFAT controls gene expression during remodeling and is activated by growth factors or histamine [109].<br>• NFAT transcription factors are involved in many normal body processes as well as in development of several diseases, such as **Kawasaki disease, inflammatory bowel diseases** and **several types of cancer**. NFAT is also being investigated as a **drug target for several different disorders** [107]. NFAT signalling is a pharmacological **target for the induction of immunosuppression**. Proteins of the NFAT family are $\text{Ca}^{2+}$-sensitive transcription factors, which are involved in **hypertrophic cardiovascular remodeling** [109]. NFAT signaling promoted |  |
| the **inflammation of wounds** [108].<br>Calcineurin-NFAT inhibitor, cyclosporine A (CsA), is the most famous **immunosuppressive drug after organ transplantations** [108]. |  |
| GP (Glycogen phosphorylase)<br>glycogenolysis pathway [109] / Glycogen phosphorylase catalyzes the rate-limiting step in glycogenolysis in animals by releasing glucose-1-phosphate from the terminal alpha-1,4-glycosidic bond. Glycogen phosphorylase is also studied as a model protein regulated by both reversible phosphorylation and allosteric effects. GP is an allosteric enzyme and with seven different binding sites discovered to date, there are multiple opportunities for modulation of its activity.<br>The inhibition of glycogen phosphorylase has been proposed as one method for **treating type 2 diabetes**. Glycogen phosphorylase is an important therapeutic **target for type 2 diabetes** having a direct influence on blood glucose levels through the glycogenolysis pathway. Considerable efforts toward the design of **drug-like GP inhibitors** have taken | 1.7–170 mHz / |
| place in recent years. Many of these inhibitors<br>are natural products and their analogues. |  |

The following parameters of Ca^2+^ oscillatory dynamic pattern are important for decoding and transmission of cell signal [5]:

1. The period of Ca^2+^ oscillations 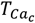 and the corresponding frequency of Ca^2+^ oscillations 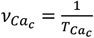. If the oscillation frequency is much lower than the typical working on-off-frequency of the specific decoding enzyme then no correct signal is transmitted [5].
2. The spike duration which is calculated as full duration half maximum (FDHM) [5]. The cumulative Ca^2+^ spike duration conveys information [98]. When the frequency is kept constant, the spike duration varies; the cumulated spike duration also changes as the cell employs pure frequency encoding [99].
3. The duty cycle. The duty cycle is the ratio of the spike duration to the period of oscillations. Maximum frequency sensitivity is observed for signals with duty cycles between 0 and 0.5 [5].
4. The amplitude of the spike [5]. The silent free calcium concentration is about 100 nmol/L in resting (non-stimulated) cells, an increase in intracellular free calcium in the form of sequence of spikes up to about 1 µmol/L is the key signal to activate the cells [100].

Synchronization index [94]. It characterizes the degree of synchronicity of calcium oscillations according to the methodology from the reference [94]. For example, individual mouse pancreatic islets exhibit oscillations in [Ca^2+^]i and insulin secretion in response to glucose *in vitro*, the oscillations of a million islets are coordinated within the human pancreas *in vivo* [94]. Islet to islet synchronization is necessary for the pancreas to produce regular pulses of insulin. Defects in this coordinating mechanism could contribute to the disrupted insulin secretion observed in Type 2 diabetes [94].

In a general case, the magnetic field frequency doesn’t coincide with the frequency of self-oscillations of the free cytosolic Ca^2+^ concentration. That is why the temporary pattern of spikes becomes quasiperiodic and additional spikes appear with amplitude increasing with increase of parameter *m*. The appearance of additional spike with infinitesimal amplitude (which increases as the magnetic field increases) is the reason of “foam-like” region in the last row of images in Fig. 3. This “foam-like” region separates the region of low WSS induced by magnetic field from the region of moderate WSS induced by magnetic field. The region of low WSS induced by magnetic field is characterized by the periodic pattern of spikes with the frequency of self-oscillations in Plank’s model. While the region of moderate WSS induced by magnetic field is characterized by quasiperiodic pattern of spikes of free intracellular Ca^2+^ concentration. The quasi-periodicity of spike’s pattern is imposed by both the frequency of self-oscillations and magnetic field frequency.

Two regimes of dynamics of concentration of free intracellular calcium are modelled: non-oscillating (the parameters for non-oscillating regime are represented in the column 4 in Table S2) and oscillating (the parameters for oscillating regime are represented in the column 5 in Table S2). The following parameters of Ca^2+^ oscillatory dynamic pattern are calculated after passing the transition process on the basis of solving the set of differential equations (7) – (10) for oscillatory regime (see the list of parameters of Plank’s model in Table S2):

- the time averaged quasi-period of oscillations of free cytosolic Ca^2+^ concentration 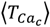and the corresponding time averaged quasi-frequency of oscillations of free cytosolic Ca2+ concentration 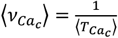. The quasi-period of quasiperiodic oscillation of intracellular calcium characterizes the time interval between the neighboring spikes of free cytosolic Ca^2+^ concentration. The quasi-period 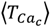 is calculated as twice of the average time interval between the neighboring maxima and minima in time-dependent solution of the differential equations (7)-(10). The initial transition process is excluded from averaging.
- the time averaged amplitude of the spike of free cytosolic Ca^2+^ concentration 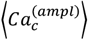. The average amplitude of spike of intracellular free cytosolic Ca^2+^ concentration is calculated as the half of the time averaged absolute value of the difference between calcium concentration in maxima (minima). The amplitudes of at least 20 maxima and minima are averaged for oscillatory regime. The initial transition process is excluded from averaging.
- the maximum amplitude of the spike of free cytosolic Ca^2+^ concentration 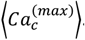. The initial transition process is excluded from calculation of the maximum amplitude.

All the calculations were carried out for over 20 quasi-periods of oscillation of the dynamic variable for oscillatory regime (see the list of parameters of Plank’s model in Table S2). Ten initial quasi-periods of oscillation of dynamic variable are excluded from calculations to exclude the transition processes passing before the oscillating regime approaches the stable periodic orbit. The amplitudes of excluded initial periods are sensitive to the initial conditions for the dynamic variables. Such approach results in the amplitudes and frequencies that are not sensitive to the initial conditions. The notation 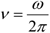 is used to define the frequency of oscillation of external magnetic field.

The exclusion of the transition process from the analysis of free calcium concentration dynamics in oscillating regime helps to exclude from the analysis the data that are very sensitive to the initial conditions for the dynamic variables of the model. Thus, the exclusion of the transition process from the analysis of free calcium concentration dynamics helps to analyze stable regimes of dynamics free calcium concentration which can be settled due to long-lasting magnetic field influence. That is why the results of the paper don’t include the analysis of the influence of single short magnetic field impulse on the dynamics of free calcium concentration. In other words, there is a spike, which we manually cut off using the code written on python programming language, excluding the time range from t = 0 to the duration of the transition process. Then self-oscillations are set if the model parameters are taken for oscillatory regime.

The following parameters of Ca^2+^ non-oscillatory dynamic pattern (see the list of parameters of Plank’s model in Table S2) are calculated on the basis of solving the set of differential equations (7) – (10):

- the time averaged intracellular free calcium concentration ⟨*Ca*_*c*_⟩. The initial transition process is not excluded from averaging.
- the time averaged standard deviation of the free calcium concentration for non-oscillatory regime 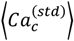. The initial transition process is not excluded from averaging; the transition process characterized by the initial spike of calcium concentration.

### Computational simulations

The results of numeric solving of equations (7-10) of intracellular calcium dynamics were carried out using python programming language, numpy and scipy packages.

### Bioinformatic analysis

The metabolic pathways of calcium decoding proteins in endothelial cells, diseases related to dysregulation of these pathway and drug targets were analyzed because magnetic field can control both the frequency and amplitude modulation of calcium signal. Search tools in bioinformatic databases (KEGG) and literature data mining were used for this purpose.

## Acknowledgements

The project is partially funded from the Mobility Program budget of the Czech Academy of Sciences and Chinese Academy of Sciences (CAS-23-01). The authors would like to thank Prof. Xin Zhang and Prof. Timothy Mitchison for the fruitful discussions.

## Code availability

The numerical codes used in this study have been deposited in a database under the accession link ….**to be added**. Additional codes are available from the corresponding authors upon request.

## Notes

See Supplemental Material at [URL will be inserted by publisher] for the open access python code as .zip file containing the python project *MagniCa*.

## Author contributions

O.G. problem statement, analytical calculations and numeric simulation, writing python code, literature data mining, developing conclusions; S.G. problem statement, numeric simulation, writing python code, literature data mining, developing conclusions; T.P. analytical calculations, literature data mining, developing conclusions; V.Z. problem statement, analytical calculations, literature data mining, developing conclusions.

## References

[1] L. Xiong and A. Garfinkel, Are Physiological Oscillations “Physiological”?, ArXiv q-bio.TO, 1 (2023).

[2] O. Garaschuk, J. Linn, J. Eilers, and A. Konnerth, Large-Scale Oscillatory Calcium Waves in the Immature Cortex, Nat. Neurosci. 3, 452 (2000).

[3] W.-G. Choi, R. Hilleary, S. J. Swanson, S.-H. Kim, and S. Gilroy, Rapid, Long-Distance Electrical and Calcium Signaling in Plants, Annu. Rev. Plant Biol. 67, 287 (2016).

[4] W. Tian, C. Wang, Q. Gao, L. Li, and S. Luan, Calcium Spikes, Waves and Oscillations in Plant Development and Biotic Interactions, Nat. Plants 6, 750 (2020).

[5] E. Smedler and P. Uhlén, Frequency Decoding of Calcium Oscillations, Biochim. Biophys. Acta - Gen. Subj. 1840, 964 (2014).

[6] M. Whitaker, Calcium at Fertilization and in Early Development, Physiol. Rev. 86, 25 (2006).

[7] C. Coburn, E. Allman, P. Mahanti, A. Benedetto, F. Cabreiro, Z. Pincus, F. Matthijssens, C. Araiz, A. Mandel, M. Vlachos, S.-A. Edwards, G. Fischer, A. Davidson, R. E. Pryor, A. Stevens, F. J. Slack, N. Tavernarakis, B. P. Braeckman, F. C. Schroeder, K. Nehrke, and D. Gems, Anthranilate Fluorescence Marks a Calcium-Propagated Necrotic Wave That Promotes Organismal Death in C. Elegans, PLoS Biol. 11, e1001613 (2013).

[8] P. De Koninck and H. Schulman, Sensitivity of CaM Kinase II to the Frequency of Ca 2+ Oscillations, Science (80-.). 279, 227 (1998).

[9] M. Colella, F. Grisan, V. Robert, J. D. Turner, A. P. Thomas, and T. Pozzan, Ca 2+ Oscillation Frequency Decoding in Cardiac Cell Hypertrophy: Role of Calcineurin/NFAT as Ca 2+ Signal Integrators, Proc. Natl. Acad. Sci. 105, 2859 (2008).

[10] G. Buzsáki and A. Draguhn, Neuronal Oscillations in Cortical Networks, Science (80-.). 304, 1926 (2004).

[11] D. Jokisch and O. Jensen, Modulation of Gamma and Alpha Activity during a Working Memory Task Engaging the Dorsal or Ventral Stream, J. Neurosci. 27, 3244 (2007).

[12] M. Rubio Ayala, T. Syrovets, S. Hafner, V. Zablotskii, A. Dejneka, and T. Simmet, Spatiotemporal Magnetic Fields Enhance Cytosolic Ca 2+ Levels and Induce Actin Polymerization via Activation of Voltage-Gated Sodium Channels in Skeletal Muscle Cells, Biomaterials 163, 174 (2018).

[13] J. J. Carson, F. S. Prato, D. J. Drost, L. D. Diesbourg, and S. J. Dixon, Time-Varying Magnetic Fields Increase Cytosolic Free Ca2+ in HL-60 Cells, Am. J. Physiol. Physiol. 259, C687 (1990).

[14] H. Wu, C. Li, M. Masood, Z. Zhang, E. González-Almela, A. Castells-Garcia, G. Zou, X. Xu, L. Wang, G. Zhao, S. Yu, P. Zhu, B. Wang, D. Qin, and J. Liu, Static Magnetic Fields Regulate T-Type Calcium Ion Channels and Mediate Mesenchymal Stem Cells Proliferation, Cells 11, 2460 (2022).

[15] T. Ikehara, H. Nishisako, Y. Minami, H. Ichinose(Sasaki), T. Shiraishi, M. Kitamura, M. Shono, H. Houchi, K. Kawazoe, K. Minakuchi, K. Yoshizaki, Y. Kinouchi, and H. Miyamoto, Effects of Exposure to a Time-Varying 1.5 T Magnetic Field on the Neurotransmitter-Activated Increase in Intracellular Ca2+ in Relation to Actin Fiber and Mitochondrial Functions in Bovine Adrenal Chromaffin Cells, Biochim. Biophys. Acta - Gen. Subj. 1800, 1221 (2010).

[16] A. Heinrich, A. Szostek, F. Nees, P. Meyer, W. Semmler, and H. Flor, Effects of Static Magnetic Fields on Cognition, Vital Signs, and Sensory Perception: A Meta-Analysis, J. Magn. Reson. Imaging 34, 758 (2011).

[17] E. A. Navarro, C. Gomez-Perretta, and F. Montes, Low Intensity Magnetic Field Influences Short-Term Memory: A Study in a Group of Healthy Students, Bioelectromagnetics 37, 37 (2016).

[18] E. A. Navarro and E. Navarro-Modesto, A Mathematical Model and Experimental Procedure to Analyze the Cognitive Effects of Audio Frequency Magnetic Fields, Front. Hum. Neurosci. 17, (2023).

[19] J. M. Henley and K. A. Wilkinson, Synaptic AMPA Receptor Composition in Development, Plasticity and Disease, Nat. Rev. Neurosci. 17, 337 (2016).

[20] D. M. Bers, Cardiac Excitation–Contraction Coupling, Nature 415, 198 (2002).

[21] J. Xu, K. Liu, T. Chen, T. Zhan, Z. Ouyang, Y. Wang, W. Liu, X. Zhang, Y. Sun, G. Xu, and X. Wang, Rotating Magnetic Field Delays Human Umbilical Vein Endothelial Cell Aging and Prolongs the Lifespan of Caenorhabditis Elegans, Aging (Albany. NY). 11, 10385 (2019).

[22] G. R. Monteith, F. M. Davis, and S. J. Roberts-Thomson, Calcium Channels and Pumps in Cancer: Changes and Consequences, J. Biol. Chem. 287, 31666 (2012).

[23] S. Feske, Calcium Signalling in Lymphocyte Activation and Disease, Nat. Rev. Immunol. 7, 690 (2007).

[24] European Bioinformatics Institute, ChEMBL Database, https://www.ebi.ac.uk/chembl/.

[25] C. J. Hutchings, P. Colussi, and T. G. Clark, Ion Channels as Therapeutic Antibody Targets, MAbs 11, 265 (2019).

[26] J. Montnach, L. A. Blömer, L. Lopez, L. Filipis, H. Meudal, A. Lafoux, S. Nicolas, D. Chu, C. Caumes, R. Béroud, C. Jopling, F. Bosmans, C. Huchet, C. Landon, M. Canepari, and M. De Waard, In Vivo Spatiotemporal Control of Voltage-Gated Ion Channels by Using Photoactivatable Peptidic Toxins, Nat. Commun. 13, 417 (2022).

[27] Z. Zhu, Z. Deng, Q. Wang, Y. Wang, D. Zhang, R. Xu, L. Guo, and H. Wen, Simulation and Machine Learning Methods for Ion-Channel Structure Determination, Mechanistic Studies and Drug Design, Front. Pharmacol. 13, (2022).

[28] M. J. Plank, D. J. N. Wall, and T. David, Atherosclerosis and Calcium Signalling in Endothelial Cells, Prog. Biophys. Mol. Biol. 91, 287 (2006).

[29] T. F. Wiesner, B. C. Berk, and R. M. Nerem, A Mathematical Model of the Cytosolic-Free Calcium Response in Endothelial Cells to Fluid Shear Stress, Proc. Natl. Acad. Sci. 94, 3726 (1997).

[30] R. O. Dull and P. F. Davies, Flow Modulation of Agonist (ATP)-Response (Ca2+) Coupling in Vascular Endothelial Cells, Am. J. Physiol. Circ. Physiol. 261, H149 (1991).

[31] J. Shen, F. W. Luscinskas, A. Connolly, C. F. Dewey, and M. A. Gimbrone, Fluid Shear Stress Modulates Cytosolic Free Calcium in Vascular Endothelial Cells, Am. J. Physiol. Physiol. 262, C384 (1992).

[32] K. Yamamoto, R. Korenaga, A. Kamiya, and J. Ando, Fluid Shear Stress Activates Ca 2+ Influx Into Human Endothelial Cells via P2X4 Purinoceptors, Circ. Res. 87, 385 (2000).

[33] R. Jacob, Calcium Oscillations in Endothelial Cells, Cell Calcium 12, 127 (1991).

[34] Y. Yokota, H. Nakajima, Y. Wakayama, A. Muto, K. Kawakami, S. Fukuhara, and N. Mochizuki, Endothelial Ca2+ Oscillations Reflect VEGFR Signaling-Regulated Angiogenic Capacity in Vivo, Elife 4, (2015).

[35] J. L. Kirschvink, A. Kobayashi-Kirschvink, and B. J. Woodford, Magnetite Biomineralization in the Human Brain., Proc. Natl. Acad. Sci. 89, 7683 (1992).

[36] F. Brem, A. M. Hirt, M. Winklhofer, K. Frei, Y. Yonekawa, H.-G. Wieser, and J. Dobson, Magnetic Iron Compounds in the Human Brain: A Comparison of Tumour and Hippocampal Tissue, J. R. Soc. Interface 3, 833 (2006).

[37] P. P. Grassi-Schultheiss, F. Heller, and J. Dobson, Analysis of Magnetic Material in the Human Heart, Spleen and Liver, BioMetals 12, 67 (1997).

[38] R. Robin Baker, J. G. Mather, and J. H. Kennaugh, Magnetic Bones in Human Sinuses, Nature 301, 78 (1983).

[39] J. L. Kirschvink, Ferromagnetic Crystals (Magnetite?) In Human Tissue., J. Exp. Biol. (1981).

[40] S. V. Gorobets, O. V. Medviediev, O. Y. Gorobets, and A. Ivanchenko, Biogenic Magnetic Nanoparticles in Human Organs and Tissues, Prog. Biophys. Mol. Biol. 135, (2018).

[41] O. Gorobets, S. Gorobets, and M. Koralewski, Physiological Origin of Biogenic Magnetic Nanoparticles in Health and Disease: From Bacteria to Humans, Int. J. Nanomedicine 12, 4371 (2017).

[42] S. Gorobets, O. Gorobets, Y. Gorobets, and M. Bulaievska, Chain-Like Structures of Biogenic and Nonbiogenic Magnetic Nanoparticles in Vascular Tissues, Bioelectromagnetics 43, 119 (2022).

[43] Y. A. Darmenko, O. Y. Gorobets, S. V. Gorobets, I. V. Sharay, and O. M. Lazarenko, Detection of Biogenic Magnetic Nanoparticles in Human Aortic Aneurysms, Acta Phys. Pol. A 133, 738 (2018).

[44] F. Brem, A. M. Hirt, C. Simon, H. G. Wieser, and J. Dobson, Characterization of Iron Compounds in Tumour Tissue from Temporal Lobe Epilepsy Patients Using Low Temperature Magnetic Methods, BioMetals (2005).

[45] O. Gorobets, S. Gorobets, I. Sharai, T. Polyakova, and V. Zablotskii, Interaction of Magnetic Fields with Biogenic Magnetic Nanoparticles on Cell Membranes: Physiological Consequences for Organisms in Health and Disease, Bioelectrochemistry 151, 108390 (2023).

[46] C. Kelly, R. K. Couch, V. T. Ha, C. M. Bodart, J. Wu, S. Huff, N. T. Herrel, H. D. Kim, A. O. Zimmermann, J. Shattuck, Y.-C. Pan, M. Kaeberlein, and A. S. Grillo, Iron Status Influences Mitochondrial Disease Progression in Complex I-Deficient Mice, Elife 12, (2023).

[47] R. Gieré, Magnetite in the Human Body: Biogenic vs. Anthropogenic, Proceedings of the National Academy of Sciences of the United States of America.

[48] B. A. Maher, Airborne Magnetite- and Iron-Rich Pollution Nanoparticles: Potential Neurotoxicants and Environmental Risk Factors for Neurodegenerative Disease, Including Alzheimer’s Disease, J. Alzheimer’s Dis. 71, 361 (2019).

[49] S. A. Winkler, F. Schmitt, H. Landes, J. de Bever, T. Wade, A. Alejski, and B. K. Rutt, Gradient and Shim Technologies for Ultra High Field MRI, Neuroimage 168, 59 (2018).

[50] A. Avasthi, C. Caro, E. Pozo-Torres, M. P. Leal, and M. L. García-Martín, Magnetic Nanoparticles as MRI Contrast Agents, Top. Curr. Chem. 378, 40 (2020).

[51] J. L. Kirschvink, A. Kobayashi-Kirschvink, and B. J. Woodford, Magnetite Biomineralization in the Human Brain., Proc. Natl. Acad. Sci. 89, 7683 (1992).

[52] A. Van de Walle, A. Plan Sangnier, A. Abou-Hassan, A. Curcio, M. Hémadi, N. Menguy, Y. Lalatonne, N. Luciani, and C. Wilhelm, Biosynthesis of Magnetic Nanoparticles from Nano-Degradation Products Revealed in Human Stem Cells, Proc. Natl. Acad. Sci. 116, 4044 (2019).

[53] S. V. Gorobets and O. Y. Gorobets, Functions of Biogenic Magnetic Nanoparticles in Organisms, Funct. Mater. 19, 18 (2012).

[54] S. Gorobets, O. Gorobets, Y. Gorobets, and M. Bulaievska, Chain-Like Structures of Biogenic and Nonbiogenic Magnetic Nanoparticles in Vascular Tissues, Bioelectromagnetics 43, 119 (2022).

[55] O. Medviediev, O. Y. Gorobets, S. V. Gorobets, and V. S. Yadrykhins’Ky, The Prediction of Biogenic Magnetic Nanoparticles Biomineralization in Human Tissues and Organs, in Journal of Physics: Conference Series, Vol. 903 (2017).

[56] D. Hautot, Q. A. Pankhurst, C. M. Morris, A. Curtis, J. Burn, and J. Dobson, Preliminary Observation of Elevated Levels of Nanocrystalline Iron Oxide in the Basal Ganglia of Neuroferritinopathy Patients, Biochim. Biophys. Acta - Mol. Basis Dis. 1772, 21 (2007).

[57] A. Kobayashi, N. Yamamoto, and J. Kirschvink, Studies of Inorganic Crystals in Biological Tissue: Magnetite in Human Tumor, Funtai Oyobi Fummatsu Yakin/Journal Japan Soc. Powder Powder Metall. 44, 294 (1997).

[58] J. Dobson, Nanoscale Biogenic Iron Oxides and Neurodegenerative Disease, 496, 1 (2001).

[59] M. Vainshtein, N. Suzina, E. Kudryashova, and E. Ariskina, New Magnet-Sensitive Structures in Bacterial and Archaeal Cells, 94, 29 (2002).

[60] O. Y. Gorobets, S. V. Gorobets, and L. V. Sorokina, Biomineralization and Synthesis of Biogenic Magnetic Nanoparticles and Magnetosensitive Inclusions in Microorganisms and Fungi, Funct. Mater. 21, 373 (2014).

[61] Y. Gorobets, S. Gorobets, O. Gorobets, A. Magerman, and I. Sharai, Biogenic and Anthropogenic Magnetic Nanoparticles in the Ploem Sieve Tubes of Plants, J. Microbiol. Biotechnol. Food Sci. e5484, (2023).

[62] S. Gorobets, O. Gorobets, I. . Sharay, and L. Yevzhyk, The Influence of Artificial and Biogenic Magnetic Nanoparticles on the Metabolism of Fungi, Funct. Mater. 28, 315 (2021).

[63] S. V. Gorobets, O. Y. Gorobets, O. V. Medviediev, V. O. Golub, and L. V. Kuzminykh, Biogenic Magnetic Nanoparticles in Lung, Heart and Liver, Funct. Mater. 24, 405 (2017).

[64] O. Veiseh, J. W. Gunn, and M. Zhang, Design and Fabrication of Magnetic Nanoparticles for Targeted Drug Delivery and Imaging ☆, Adv. Drug Deliv. Rev. 62, 284 (2010).

[65] J. Chomoucka, J. Drbohlavova, D. Huska, V. Adam, R. Kizek, and J. Hubalek, Magnetic Nanoparticles and Targeted Drug Delivering, Pharmacol. Res. 62, 144 (2010).

[66] P. M. Price, W. E. Mahmoud, A. A. Al-Ghamdi, and L. M. Bronstein, Magnetic Drug Delivery: Where the Field Is Going, Front. Chem. 6, 1 (2018).

[67] S. M. Mirvakili and R. Langer, Wireless On-Demand Drug Delivery, Nat. Electron. 4, 464 (2021).

[68] S. V. Gorobets, O. Y. Gorobets, Y. M. Chyzh, and D. V. Sivenok, Magnetic Dipole Interaction of Endogenous Magnetic Nanoparticles with Magnetoliposomes for Targeted Drug Delivery, Biophys. (Russian Fed. 58, 379 (2013).

[69] H. Gavilán, S. K. Avugadda, T. Fernández-Cabada, N. Soni, M. Cassani, B. T. Mai, R. Chantrell, and T. Pellegrino, Magnetic Nanoparticles and Clusters for Magnetic Hyperthermia: Optimizing Their Heat Performance and Developing Combinatorial Therapies to Tackle Cancer, Chem. Soc. Rev. 50, 11614 (2021).

[70] A. Rajan and N. K. Sahu, Review on Magnetic Nanoparticle-Mediated Hyperthermia for Cancer Therapy, J. Nanoparticle Res. 22, 319 (2020).

[71] J. L. Kirschvink, M. M. Walker, S.-B. Chang, A. E. Dizon, and K. A. Peterson, Chains of Single-Domain Magnetite Particles in Chinook Salmon,Oncorhynchus Tshawytscha, J. Comp. Physiol. A 157, 375 (1985).

[72] J. Kirschvink, Magnetite-Based Magnetoreception, Curr. Opin. Neurobiol. 11, 462 (2001).

[73] E. M. Alfsen, F. C. Størmer, A. Njå, and L. Walløe, A Proposed Tandem Mechanism for Memory Storage in Neurons Involving Magnetite and Prions, Med. Hypotheses 119, 98 (2018).

[74] S. Miclaus, C. Iftode, and A. Miclaus, WOULD THE HUMAN BRAIN BE ABLE TO ERECT SPECIFIC EFFECTS DUE TO THE MAGNETIC FIELD COMPONENT OF AN UHF FIELD VIA MAGNETITE NANOPARTICLES?, Prog. Electromagn. Res. M 69, 23 (2018).

[75] I. Miller and B. Lonetree, The Sedona Effect: Correlations between Geomagnetic Anomalies, EEG Brainwaves & Schumann Resonance, J. Conscious. Explor. Res. 4, 630 (2013).

[76] A. Kobayashi, N. Yamamoto, and J. Kirschvink, Studies of Inorganic Crystals in Biological Tissue: Magnetic in Human Tumor., J. Japan Soc. Powder Powder Metall. 44, 294 (1997).

[77] O. Y. Gorobets, S. V. Gorobets, and Y. I. Gorobets, Biogenic Magnetic Nanoparticles. Biomineralization in Prokaryotes and Eukaryotes, in In Dekker Encyclopedia of Nanoscience and Nanotechnology, Third Edition. CRC Press: New York (2014), pp. 300–308.

[78] J. W. Putney, L. M. Broad, F.-J. Braun, J.-P. Lievremont, and G. S. J. Bird, Mechanisms of Capacitative Calcium Entry, J. Cell Sci. 114, 2223 (2001).

[79] E. Roux, P. Bougaran, P. Dufourcq, and T. Couffinhal, Fluid Shear Stress Sensing by the Endothelial Layer, Front. Physiol. 11, (2020).

[80] R. Skalak, A. Tozeren, R. P. Zarda, and S. Chien, Strain Energy Function of Red Blood Cell Membranes, Biophys. J. 13, 245 (1973).

[81] J. J. Lacroix, W. M. Botello-Smith, and Y. Luo, Probing the Gating Mechanism of the Mechanosensitive Channel Piezo1 with the Small Molecule Yoda1, Nat. Commun. 9, 2029 (2018).

[82] J.-J. Chiu and S. Chien, Effects of Disturbed Flow on Vascular Endothelium: Pathophysiological Basis and Clinical Perspectives, Physiol. Rev. 91, 327 (2011).

[83] S. Sonam, L. Balasubramaniam, S.-Z. Lin, Y. M. Y. Ivan, I. Pi-Jaumà, C. Jebane, M. Karnat, Y. Toyama, P. Marcq, J. Prost, R.-M. Mège, J.-F. Rupprecht, and B. Ladoux, Mechanical Stress Driven by Rigidity Sensing Governs Epithelial Stability, Nat. Phys. 19, 132 (2023).

[84] L. Cheng, J. Li, H. Sun, and H. Jiang, Appropriate Mechanical Confinement Inhibits Multipolar Cell Division via Pole-Cortex Interaction, Phys. Rev. X 13, 011036 (2023).

[85] L. F. Jaffe, Fast Calcium Waves, Cell Calcium 48, 102 (2010).

[86] J. Galvanovskis, J. Sandblom, B. Bergqvist, S. Galt, and Y. Hamnerius, Cytoplasmic Ca2+ Oscillations in Human Leukemia T-Cells Are Reduced by 50 Hz Magnetic Fields, Bioelectromagnetics 20, 269 (1999).

[87] C. X. Wang, I. A. Hilburn, D.-A. Wu, Y. Mizuhara, C. P. Cousté, J. N. H. Abrahams, S. E. Bernstein, A. Matani, S. Shimojo, and J. L. Kirschvink, Transduction of the Geomagnetic Field as Evidenced from Alpha-Band Activity in the Human Brain, Eneuro 6, ENEURO.0483 (2019).

[88] Y. Kasai, T. Yamazawa, T. Sakurai, Y. Taketani, and M. Iino, Endothelium-Dependent Frequency Modulation of Ca 2+ Signalling in Individual Vascular Smooth Muscle Cells of the Rat, J. Physiol. 504, 349 (1997).

[89] Moccia, Negri, Shekha, Faris, and Guerra, Endothelial Ca2+ Signaling, Angiogenesis and Vasculogenesis: Just What It Takes to Make a Blood Vessel, Int. J. Mol. Sci. 20, 3962 (2019).

[90] Simonetti and Mohaupt, Kalzium Und Blutdruck, Ther. Umschau 64, 249 (2007).

[91] F. E. Curry, Modulation of Venular Microvessel Permeability by Calcium Influx into Endothelial Cells, FASEB J. 6, 2456 (1992).

[92] P. J. Dalal, D. P. Sullivan, E. W. Weber, D. B. Sacks, M. Gunzer, I. M. Grumbach, J. Heller Brown, and W. A. Muller, Spatiotemporal Restriction of Endothelial Cell Calcium Signaling Is Required during Leukocyte Transmigration, J. Exp. Med. 218, (2021).

[93] G. Guerra, A. Lucariello, A. Perna, L. Botta, A. De Luca, and F. Moccia, The Role of Endothelial Ca2+ Signaling in Neurovascular Coupling: A View from the Lumen, Int. J. Mol. Sci. 19, 938 (2018).

[94] M. Zhang, B. Fendler, B. Peercy, P. Goel, R. Bertram, A. Sherman, and L. Satin, Long Lasting Synchronization of Calcium Oscillations by Cholinergic Stimulation in Isolated Pancreatic Islets, Biophys. J. 95, 4676 (2008).

[95] S. Ghodbane, A. Lahbib, M. Sakly, and H. Abdelmelek, Bioeffects of Static Magnetic Fields: Oxidative Stress, Genotoxic Effects, and Cancer Studies, Biomed Res. Int. 2013, 1 (2013).

[96] A. D. Rosen, Mechanism of Action of Moderate-Intensity Static Magnetic Fields on Biological Systems, Cell Biochem. Biophys. 39, 163 (2003).

[97] H. Lei, Y. Pan, R. Wu, and Y. Lv, Innate Immune Regulation Under Magnetic Fields With Possible Mechanisms and Therapeutic Applications, Front. Immunol. 11, (2020).

[98] S. Tóth, D. Huneau, B. Banrezes, and J. P. Ozil, Egg Activation Is the Result of Calcium Signal Summation in the Mouse, Reproduction 131, (2006).

[99] C. Salazar, A. Z. Politi, and T. Höfer, Decoding of Calcium Oscillations by Phosphorylation Cycles: Analytic Results, Biophys. J. 94, (2008).

[100] F. Moccia, Update on Vascular Endothelial Ca 2+ Signalling: A Tale of Ion Channels, Pumps and Transporters, World J. Biol. Chem. 3, 127 (2012).

[101] W. Zhang and H. T. Liu, MAPK Signal Pathways in the Regulation of Cell Proliferation in Mammalian Cells, Cell Res. 12, 9 (2002).

[102] R. J. Roth Flach, C.-A. Guo, L. V. Danai, J. C. Yawe, S. Gujja, Y. J. K. Edwards, and M. P. Czech, Endothelial Mitogen-Activated Protein Kinase Kinase Kinase Kinase 4 Is Critical for Lymphatic Vascular Development and Function, Mol. Cell. Biol. 36, 1740 (2016).

[103] S. Kempe, NF-B Controls the Global pro-Inflammatory Response in Endothelial Cells: Evidence for the Regulation of a pro-Atherogenic Program, Nucleic Acids Res. 33, 5308 (2005).

[104] B. Hoesel and J. A. Schmid, The Complexity of NF-?B Signaling in Inflammation and Cancer, Mol. Cancer 12, 86 (2013).

[105] T. Lawrence, The Nuclear Factor NF-B Pathway in Inflammation, Cold Spring Harb. Perspect. Biol. 1, a001651 (2009).

[106] T. Liu, L. Zhang, D. Joo, and S.-C. Sun, NF-?B Signaling in Inflammation, Signal Transduct. Target. Ther. 2, 17023 (2017).

[107] G. R. Crabtree and E. N. Olson, NFAT Signaling, Cell 109, S67 (2002).

[108] T. Manabe, H. Park, and T. Minami, Calcineurin-Nuclear Factor for Activated T Cells (NFAT) Signaling in Pathophysiology of Wound Healing, Inflamm. Regen. 41, 26 (2021).

[109] A. Rinne, K. Banach, and L. A. Blatter, Regulation of Nuclear Factor of Activated T Cells (NFAT) in Vascular Endothelial Cells, J. Mol. Cell. Cardiol. 47, 400 (2009).

