## Supplementary Material for "Modulation of Calcium Signaling and Metabolic Pathways in Endothelial Cells with Magnetic Fields"

**Supplementary material. Magnetic shear stress in the modified Plank's model $Ca^{2+}$ dynamics in endothelial cells taking on account magnetic field influence on the chains of magnetic nanoparticles embedded in cell membrane**

The transformation from dimensional dynamic variables to dimensionless ones is given in Table S1.

Table S1. Dynamic variables in the Plank's model for calcium dynamics.

| Dynamic variable | Dimensional notation | Dimensionless notation |
|---|---|---|
| Concentrations of IP3 in the cytosol | $i$ | $i' = \frac{i}{K_4}$ |
| Free $Ca^{2+}$ in the cytosol | $Ca_c, \mu M$ | $Ca'_c = \frac{Ca_c}{K_4}$ |
| Buffered $Ca^{2+}$ in the cytosol | $Ca_b, \mu M$ | $Ca'_b = \frac{Ca_b}{K_4}$ |
| $Ca^{2+}$ in the internal stores | $Ca_s, \mu M$ | $Ca'_s = \frac{Ca_s}{K_4}$ |
| Sum of free $Ca^{2+}$ in the cytosol and buffered $Ca^{2+}$ in the cytosol | $Ca_t = Ca_c + Ca_b$, $\mu M$ | $Ca'_t = \frac{Ca_t}{K_4}$ |

Let's take on account

$$2Ca_c = Ca_t - B_T - \frac{k_7}{k_6} + \left(\left(Ca_t - B_T - \frac{k_7}{k_6}\right)^2 + 4\frac{k_7}{k_6}Ca_t\right)^{1/2} \quad \text{(S1)}$$

Substituting (S1) into the set of equations (10)-(13) we obtain:

$$\frac{di}{dt} = k_1 \frac{\phi}{K_c+\phi} \cdot \frac{Ca_c}{K_1+Ca_c} - k_2 i \tag{S2}$$

$$\frac{dCa_t}{dt} = k_3 \frac{Ca_c}{K_{CICR}+Ca_c}\left(\frac{i}{K_2+i}\right)^3 Ca_s - k_4\left(\frac{Ca_C}{K_3+Ca_c}\right)^2 + k_5 Ca_s^2 + k_{CCE}\left(Ca_{s,0} - Ca_s\right)(Ca_{ex} - Ca_c) + \frac{q_{\max}}{1+\alpha\exp\left(-\frac{f_e W(\tau)}{kT_e N}\right)} - \frac{k_8}{K_4+Ca_c} \tag{S3}$$

$$\frac{dCa_s}{dt} = -V_r\left(k_3 \frac{Ca_c}{K_{CICR}+Ca_c}\left(\frac{i}{K_2+i}\right)^3 Ca_s - k_4\left(\frac{Ca_C}{K_3+Ca_c}\right)^2 + k_5 Ca_s^2\right) \tag{S4}$$

The initial conditions are used for the equations (12)-(14):

$$i(0) = 0 \tag{S5}$$

$$Ca_t(0) = Ca_0\left(\frac{Ca_0+B_T+k_7/k_6}{Ca_0+k_7/k_6}\right) \tag{S6}$$

$$Ca_s(0) = Ca_{s,0} \tag{S7}$$

The non-dimensionless equations have the form after non-dimensionalisation (Table S1, Table S2):

$$\frac{di'}{dt'} = k_1' \frac{\phi'}{K_c'+\phi'} \cdot \frac{Ca_c'}{K_1'+Ca_c'} - k_2' i' \tag{S8}$$

$$\frac{dCa_t'}{dt'} = k_3' \frac{Ca_c'}{K_{CICR}'+Ca_c'}\left(\frac{i'}{K_2+i'}\right)^3 Ca_s' - k_4'\left(\frac{Ca_c'}{K_3'+Ca_c'}\right)^2 + k_5' Ca_s'^2 + k_{CCE}'\left(Ca_{s,0}' - Ca_s'\right)(Ca_{ex}' - Ca_c') + \frac{q_{max}'}{1+\alpha\exp\left(-W'(\tau')\right)} - \frac{k_8'}{K_4'+Ca_c'} \tag{S9}$$

$$\frac{dCa_s'}{dt'} = -V_r'\left(k_3' \frac{Ca_c'}{K_{CICR}'+Ca_c'}\left(\frac{i'}{K_2+i'}\right)^3 Ca_s' - k_4'\left(\frac{Ca_c'}{K_3'+Ca_c'}\right)^2 + k_5' Ca_s'^2\right) \tag{S10}$$

$$W'(\tau') = \frac{f_e W_0'\left[\varepsilon'\tau' + \sqrt{16\delta'^2 + \varepsilon'^2\tau'^2} - 4\delta'\right]^2}{\left[\varepsilon'\tau' + \sqrt{16\delta'^2 + \varepsilon'^2\tau'^2}\right]}. \tag{S11}$$

The results of numeric solving of equations (S8)-(S11) using Python programming language, numpy and scipy packages.

Table S2. Parameters in the Plank's model for calcium dynamics modified to take on account the WSS induced by magnetic field.

| Parameter used in the model for calcium dynamics | Dimensional parameter values for non-oscillatory regime | Relation between dimensional and dimension-less parameter | Dimension-less parameter values for non-oscillatory regime | Dimension-less parameter values for oscillatory regime |
|---|---|---|---|---|
| Maximum value of spatial derivative of magnetic field flux density | $G_0 = 30\, Tm^{-1}$ | | | |
| Magnetic field flux density of the Earth | $B = 0.5 \cdot 10^{-4}\, T$ | | | |
| Biogenic or nonbiogenic magnetic nanoparticle radius | $r = 100 \cdot 10^{-9} m$ | | | |
| Magnetization of biogenic or | $M_s = 510 \cdot 10^3 Am^{-1}$ | | | |

| | | | |
|---|---|---|---|
| nonbiogenic magnetic nanoparticle | | | |
| Frequency of external gradient magnetic field oscillation | $\omega = 4 \cdot 10^{-4}\ s^{-1}$ | $\omega' = \frac{\omega}{T}$ | $\omega' = 0.2$ |
| Amplitude of magnetic WSS (WSS) induced by gradient magnetic field | $P_0 = 2.04\ Pa$ $P_0 = \frac{4rM_sG_0}{3}$ | $\tau'_{magn} = \frac{P_0}{\rho U_0^2}$ | $\tau'_{magn} =$ 18.9 for artery (Plank's model) $\tau'_{magn} =$ 906.7 for capillary |
| Amplitude of magnetic WSS (WSS) induced by uniform magnetic field | $P_0 = 1.3\ Pa$ $P_0 = \frac{\pi}{6N} M_S B \sin\theta sin\gamma$ | $\tau'_{magn} = \frac{P_0}{\rho U_0^2}$ | $\tau'_{magn} =$ 12.3 for artery (Plank's model) $\tau'_{magn} =$ 593 for capillary |

| | | | | |
|---|---|---|---|---|
| Angle between the normal to the cell membrane and magnetic field | $\theta$ | | | |
| Angle between in-plane component of magnetic field and magnetization of the chain of magnetic nanoparticles | $\gamma$ | | | |
| Concentration of ATP at the cell surface | $\phi$ [24] | $\phi' = \frac{\phi}{\phi_0}$ | $\phi' =$ 0.9 [24] | |
| Reference ATP concentration | $\phi_0 = 0.1\ \mu M$ [24] | | | |
| IP3 production rate | $k_1 = 5.46 \cdot 10^{-3}\ \mu M s^{-1}$ [24] | $k_1' = \frac{k_1 T}{K_4}$ | $k_1' =$ 8.53 [24] | $k_1' =$ 35.8 [24] |
| IP3 decay rate | $k_2 = 0.2\ s^{-1}$ [24] | $k_2' = k_2 T$ | $k_2' =$100 [24] | |

| | | | | |
|---|---|---|---|---|
| $Ca^{2+}$ release rate | $k_3 = 6.64\ s^{-1}$ [24] | $k_3' = k_3 T$ | $k_3' = 3320$ [24] | |
| $Ca^{2+}$ resequestration rate | $k_4 = 5\ \mu M s^{-1}$ [24] | $k_1' = \frac{k_4 T}{K_4}$ | $k_4' = 7810$ [24] | $k_4' = 7.81 \cdot 10^4$ [24] |
| $Ca^{2+}$ leak rate | $k_5 = 10^{-7}\ \mu M^{-1} s^{-1}$ [24] | $k_5' = k_5 T K_4$ | $k_5' = 1.6 \cdot 10^{-5}$ [24] | |
| $Ca^{2+}$ buffering rate | $k_6 = 100\ \mu M^{-1} s^{-1}$ [24] | | | |
| $Ca^{2+}$ debuffering rate | $k_7 = 300\ s^{-1}$ [24] | | | |
| Ratio of $Ca^{2+}$ buffering rates | | $k_b' = \frac{k_7 T}{k_6 K_4}$ | $k_b = 9.38$ [24] | |
| $Ca^{2+}$ efflux rate | $k_8 = 24.7\ \mu M s^{-1}$ [24] | $k_8' = \frac{k_8 T}{K_4}$ | $k_8' = 38600$ [24] | |
| Max. WSS-induced $Ca^{2+}$ influx rate | $q_{\max} = 17.6\ \mu M s^{-1}$ [24] | $q_{max}' = \frac{q_{max} T}{K_4}$ | $q_{max}' = 27500$ [24] | |
| CCE rate | $k_{CCE} = 8 \cdot 10^{-7}\ \mu M^{-1} s^{-1}$ [24] | $k_{CCE}' = k_{CCE} T K_4$ | $k_{CCE}' = 1.28 \cdot 10^{-4}$ [24] | |
| Resting cytosolic $Ca^{2+}$ | $Ca_0 = 0.1\ \mu M$ [24] | $Ca_0' = \frac{Ca_0}{K_4}$ | $Ca_0' = 0.313$ [24] | |

| | | | | |
|---|---|---|---|---|
| concentration | | | | |
| Resting stored $Ca^{2+}$ concentration | $Ca_{s,0} = 2828\,\mu M$ [24] | $Ca'_{s,0} = \frac{Ca_{s,0}}{K_4}$ | $Ca'_{s,0} =$ 8840 [24] | |
| External $Ca^{2+}$ concentration | $Ca_{ex} = 1500\,\mu M$ [24] | $Ca'_{ex} = \frac{Ca_{ex}}{K_4}$ | $Ca'_{ex} =$ 4690 [24] | |
| Concentration of $Ca^{2+}$ buffering sites | $B_T = 120\,\mu M$ [24] | $B'_T = \frac{B_T}{K_4}$ | $B'_T =$ 375 [24] | |
| Michaelis–Menten constants | $K_{CICR} = 0\,\mu M$ [24] | $K'_{CICR} = \frac{K_{CICR}}{K_4}$ | 0 [24] | |
| Michaelis–Menten constants | $K_1 = 0\,\mu M$ [24] | $K'_1 = \frac{K_1}{K_4}$ | $K'_1 = 0$ [24] | $K'_1 = 1$ [24] |
| Michaelis–Menten constants | $K_2 = 0.2\,\mu M$ [24] | $K'_2 = \frac{K_2}{K_4}$ | $K'_2 =$ 0.625 [24] | |
| Michaelis–Menten constants | $K_3 = 0.15\,\mu M$ [24] | $K'_3 = \frac{K_3}{K_4}$ | $K'_3 =$ 0.469 [24] | |
| Michaelis–Menten constants | $K_4 = 0.32\,\mu M$ [24] | | | |

| | | | |
|---|---|---|---|
| Michaelis–Menten constants | $K_c = 0.026\ \mu M$ [24] | $K_c' = \frac{K_c}{\phi_0}$ | $K_c' =$ 0.26 [24] |
| Membrane shear modulus | $\delta = 10^{-5}\ kg \cdot s^{-2}$ [24] | $\delta' = \frac{\delta}{\rho U_0^2 l}$ | $\delta' = 2.63$ for artery (Plank's model) [24] $\delta' = 126$ for capillary |
| Density of water | $\rho = 1000\ kgm^{-3}$ [24] | | |
| Reference velocity | $U_0 = 1.04 \cdot 10^{-2}\ ms^{-1}$ [24] for artery (Plank's model) $U_0 = 0.2\ ms^{-1}$ typical for artery $U_0 = 1.5 \cdot 10^{-3}\ ms^{-1}$ typical for capillary | | |
| Cell length in direction of flow | $l = 3.5 \cdot 10^{-5}\ m$ [24] | | |
| Plasma membrane load fraction | | | $\varepsilon' =$ 0.1 [24] |
| Plasma membrane energy gating fraction | | | $f_e' = 0.0134$ [24] |

| | | | |
|---|---|---|---|
| Ratio of cytosolic and ER volumes | | | $V_r' =$ 3.5 [24] |
| Strain energy density constant | | $W_0' = \frac{\rho U_0^2 l}{8kT_e N}$ | $W_0' =$ 111 [24] for artery $W_0' = 2.3$ for capillary |
| Temperature | $T_e = 310\ K$ | | |
| $Ca^{2+}$ channel area density | $N_0 = 10^{12}\ m^{-2}$ [24] | | |
| Boltzmann constant | $k = 1.3807 \cdot 10^{-23}\ kg \cdot m^2 s^{-2} K^{-1}$ | | |
| time | $t, s$ | $t' = \frac{t}{T}$ | |
| Reference timescale | $T = 500\ s$ | | |
| Characteristic WSS for ATP production | $\tau_m = 1\ kg \cdot m^{-1} s^{-2}$ [24] | | $\tau_m' =$ 9.22 [24] |
| WSS | $\tau =$ 1 Pa for artery $\tau = 0.1$ Pa [75] for capillary | $\tau' = \frac{\tau}{\rho U_0^2}$ [24] | $\tau' = 9.2$ for artery $\tau' = 44$ for capillary |

First, we reproduce the results of Plank's model for artery without magnetic field influence in non-oscillatory regime. Then we calculate the influence of oscillating magnetic shear stress. The set of dimensionless parameters for 'non-oscillatory regime', 'artery' is given in Table S2.

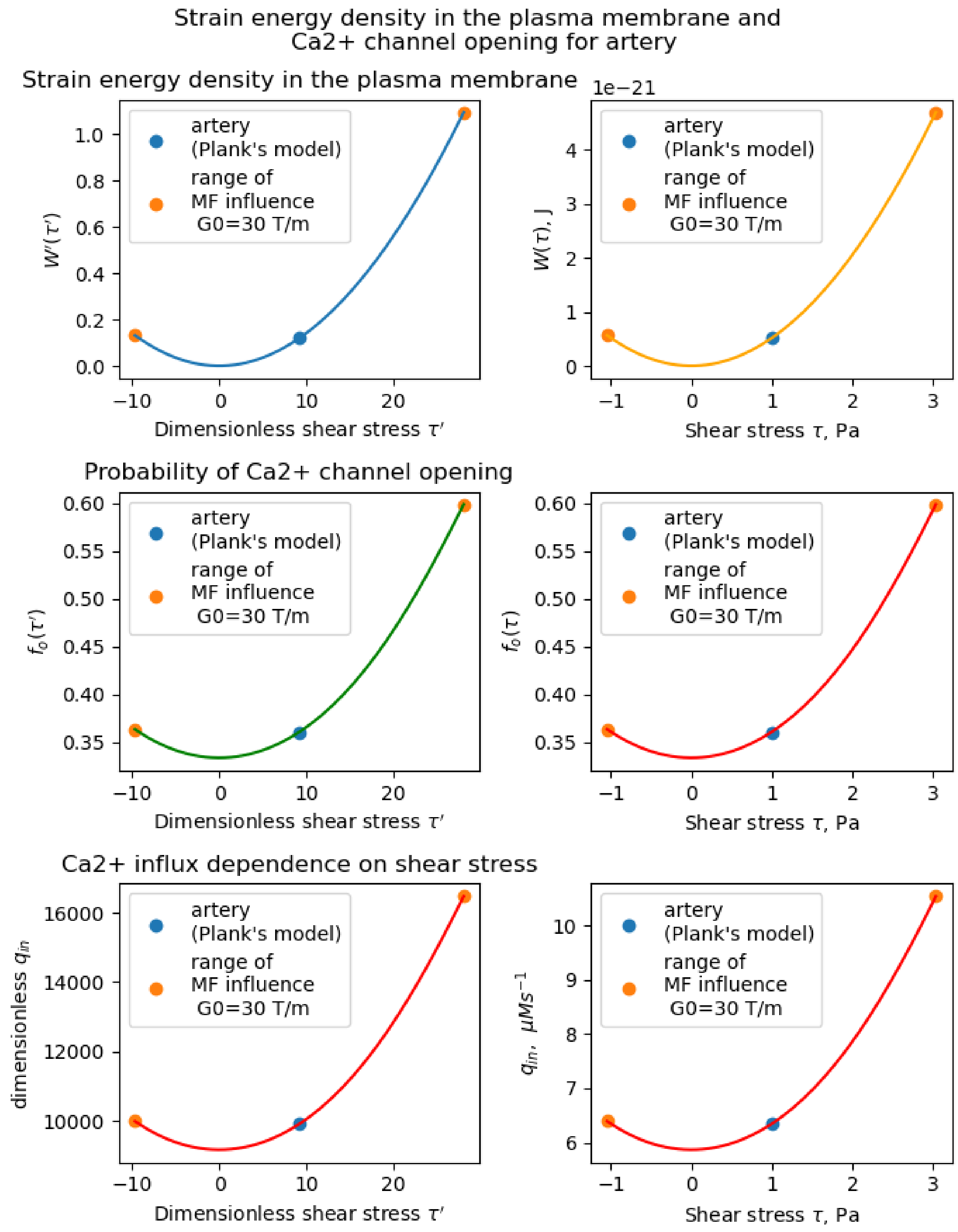

Fig. S1 Range of oscillating magnetic field influence ($G_0 \in [0,30]\frac{T}{m}$) on strain energy density in the plasma membrane and $Ca^{2+}$ channel opening for artery in Plank's model [24].

**Code availability**

The codes used in this study are available at www…. as .zip file containing the python project MagniCa.

**Additional information**

**Supplementary information**

The online version contains Supplementary Material available at www…..

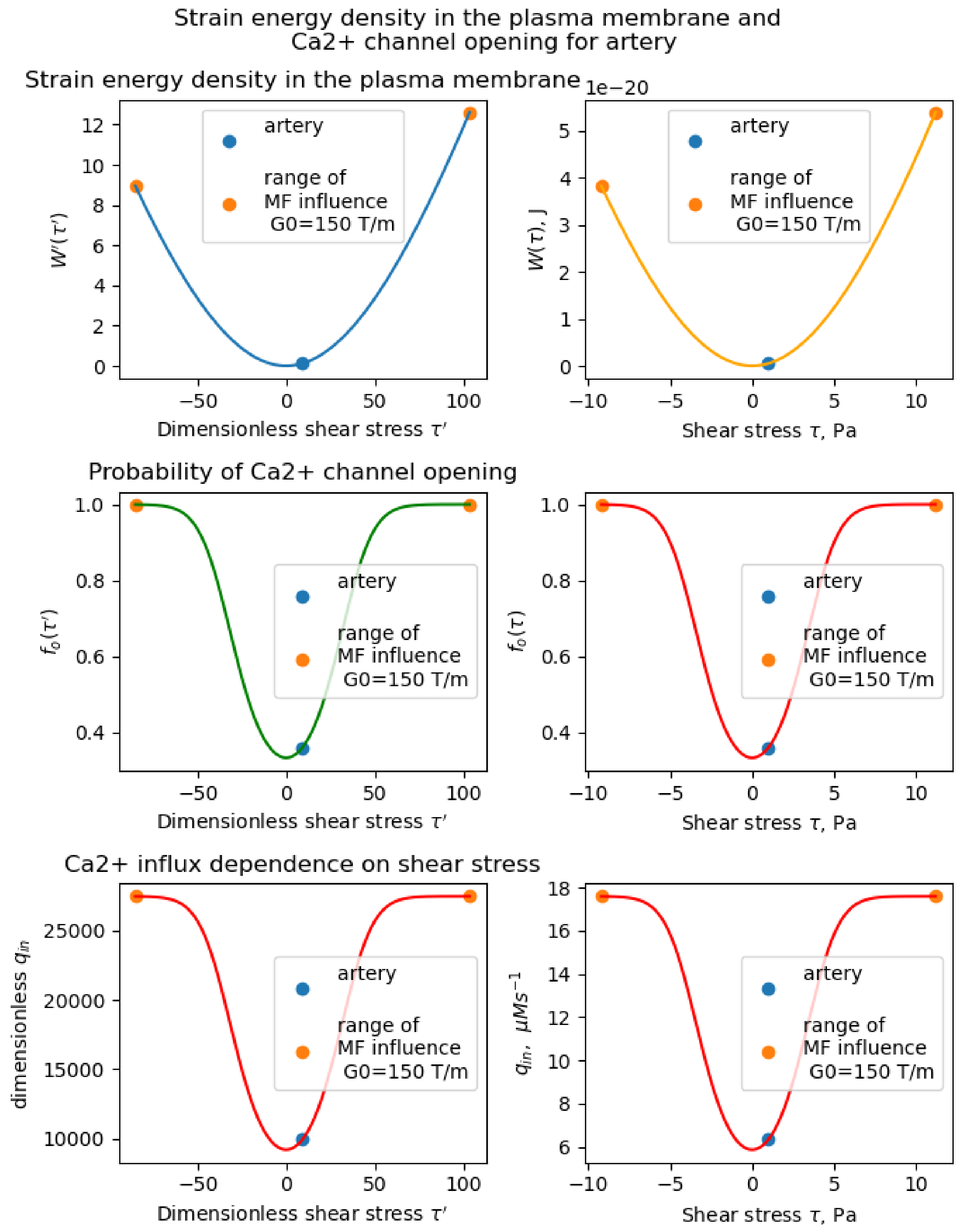

Fig. S2. Range of oscillating magnetic field influence ($G_0 \in [0{,}150]\frac{T}{m}$) on strain energy density in the plasma membrane and $Ca^{2+}$ channel opening for artery in Plank's model [24].

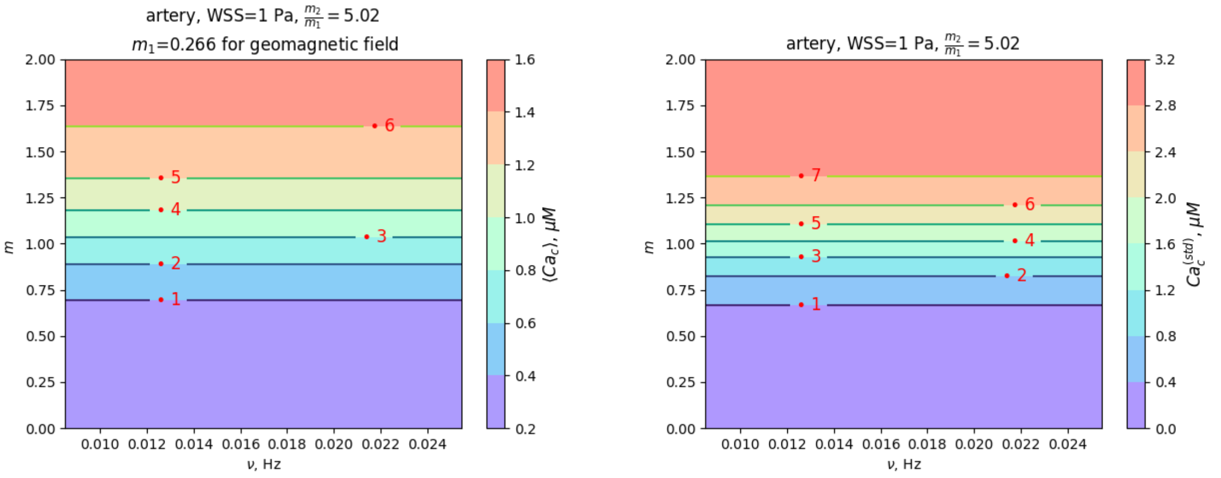

A B

Fig. S3. Colormaps visualizing magnetic field effects on dynamics of intracellular free $Ca^{2+}$ concentration for artery in non-oscillatory regime (Table S2).

WSSs created by magnetic field below "magnetic field saturation" threshold $\tau_{magn}$ ($G_0 < 80$ T/m in the case of gradient magnetic field) are much less than the normal blood pressure (1840 Pa in artery, upper limit of normal is 2660 Pa [107], 4655 Pa in arteriolar [108]; capillary pressure differs markedly among tissues in the range of 665 Pa – 6650 Pa). Therefore, the direction of the magnetic force in plane of membrane is very important for ion channel gating. The out of plane magnetic force doesn't gate the ion channels as well as the out of plane force exerted on membrane by blood pressure itself doesn't gate the mechanosensitive ion channel.

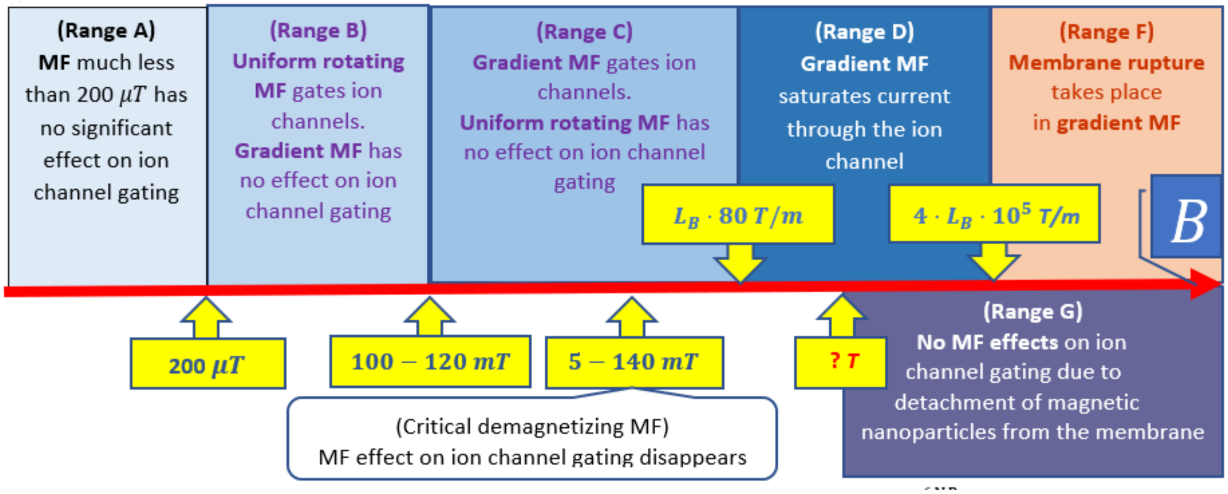

Fig. S4. **Roadmap of expected magnetic field effects. Range A:** Magnetic field (MF) $B \ll \frac{6NP_s}{\pi M_S}$ has no significant effect on ion channel gating because magnetic WSS is much less than WSS saturating $Ca^{2+}$ ion channel. **Range B:** Uniform rotating MF gates ion channels. Gradient MF has no effect on ion channel gating for $L_B$ greater than the length of chain of magnetic nanoparticles where $L_B$ is the characteristic scale of magnetic field gradient. **Range C:** Gradient MF gates ion channels. Uniform rotating MF has no effect on ion channel gating because magnetic field is equal to saturation magnetic field for magnetic nanoparticle chain and the chain is magnetized parallel to the external magnetic field $B \approx 100 - 120\ mT$. **Range D:** WSS of gradient magnetic field $B_{cr} \approx L_B G_s,\ G_s \approx 80\ T/m$ saturates $Ca^{2+}$ ion channel. **Range F:** Magnetic WSS is high resulting in membrane rupture $G_{cr} = \frac{3P_{cr}}{4rM_s} \approx 4 \cdot 10^5$ T/m, $B_{cr} = G_{cr} L_B$. **Range G:** No magnetic field effects on ion channel gating due to detachment of magnetic nanoparticles from the membrane.
